# Loneliness is linked to specific subregional alterations in hippocampus-default network co-variation

**DOI:** 10.1101/2021.08.19.456905

**Authors:** Chris Zajner, Nathan Spreng, Danilo Bzdok

## Abstract

Social interaction complexity makes humans unique. But in times of social deprivation this strength risks to expose important vulnerabilities. Human social neuroscience studies have placed a premium on the default network (DN). In contrast, hippocampus (HC) subfields have been intensely studied in rodents and monkeys. To bridge these two literatures, we here quantified how DN subregions systematically co-vary with specific HC subfields in the context of subjective social isolation (i.e., loneliness). By co-decomposition using structural brain scans of ∼40,000 UK Biobank participants, loneliness was specially linked to midline subregions in the uncovered DN patterns. These association cortex signatures coincided with concomitant HC patterns implicating especially CA1 and molecular layer. These patterns also showed a strong affiliation with the fornix white-matter tract and the nucleus accumbens. In addition, separable signatures of structural HC-DN co-variation had distinct associations with the genetic predisposition for loneliness at the population level.

## Introduction

Social isolation can be detrimental. Accumulating evidence suggests social disconnection as a major risk factor for morbidity and mortality (Bzdok and Dunbar, 2020; Holt-Lunstad et al., 2015). Yet, it is becoming increasingly clear that experiencing feelings of social isolation, namely loneliness, has distinct biological correlates compared to an objective lack of social contact with others (Cacioppo and Hawkley, 2009). Indeed, it has been found that only ∼10-20% of adults who live alone report feeling lonely (Beutel et al., 2017). Conversely, individuals who are well-surrounded by others can still experience a feeling of inadequate social connection (Hawkley and Cacioppo, 2010). Therefore, it is widely acknowledged that there can be a divide between subjective satisfaction with their social relationships and people’s objective level of social support.

Lonely individuals are particularly characterized by consistent cognitive biases related to processing social cues in the environment. Additionally, loneliness was reported to entail a state of hyper-vigilance for threats in one’s social environment (Hawkley and Cacioppo, 2010). According to some authors, an aversive feeling of loneliness serves as biological warning signal that alerts individuals to improve their social relationships (Cacioppo and Cacioppo, 2018b). Yet, the impact of loneliness extends well beyond the realm of social interaction. For example, previous studies showed that lonely individuals have reduced cognitive control (Cacioppo et al., 2000), priority for nonsocial rewards over social rewards (Cacioppo et al., 2009), reduced immune response to viruses (Cacioppo and Cacioppo, 2018b; Cole et al., 2015), heightened stress response (Cacioppo and Cacioppo, 2018a; Cacioppo et al., 2015), poorer mental health (Holt-Lunstad et al., 2015), increased susceptibility to major psychiatric disorders (Holt-Lunstad et al., 2015), as well as higher risks for alcohol abuse (Åkerlind and Hörnquist, 1992), committing suicide (Beutel et al., 2017; Holt-Lunstad et al., 2015), cardiovascular disease (Caspi et al., 2006; Hawkley et al., 2003), cognitive decline, and Alzheimer’s disease (Holwerda et al., 2014; Lara et al., 2019; Wilson et al., 2007). In fact, the hippocampus (Braak and Braak, 1996) and default network (DN) (Buckner et al., 2008) have long been recognized as a primary neural pathway implicated in the pathophysiology of Alzheimer’s disease.

The disruption of memory is one of the hallmarks of Alzheimer’s disease (Sperling et al., 2010), while the DN and HC have both been implicated in environment-independent processing, such as episodic memory and mental scene construction (Burgess et al., 2002; Dohmatob et al., 2020a; Hassabis et al., 2007; Hassabis and Maguire, 2009). That these two neural systems should relate to loneliness may be unsurprising. This is because the past literature on loneliness has insisted on the importance of rumination on self-focused memories (Zawadzki et al., 2013), poor executive control (Cacioppo et al., 2000; Campbell et al., 2006), and negative bias in the perception of social cues (cf. above). In fact, the aspects of loneliness which rely upon internally-generated dimensions of cognition have been argued to especially relate to the DN (Andrews-Hanna et al., 2014a). Additionally, in rodents and monkeys, the HC has been found to support highly synchronized and spontaneous neural activity bursts during states of rest: so-called sharp-wave ripples (SWR). These rapid electrophysiological bursts have been described to originate in specific anatomical areas of the hippocampus, such as CA1, known to be closely affiliated with DN regions, especially with the medial prefrontal cortex (mPFC) (Buzsáki, 2015). For example, in monkey brains, hippocampal CA1 neurons have been shown to send direct axonal output connections to the mPFC through the fornix white-matter pathway (Aggleton et al., 2015). This fact could in part explain the recent finding that the fornix is the major white-matter tract that is most strongly linked to loneliness (Spreng et al., 2020). The nucleus accumbens (NAc), which is another major target site of hippocampal CA1 neurons and the fornix (Aggleton et al., 2015; Friedman et al., 2002), has likewise been linked to the craving for social connection (Kiesow et al., 2020). Alteration of the NAc in loneliness, and thus the reward circuitry of the brain, may therefore be a product of this common anatomical pathway. In light of these details, it has been concluded that since “the pyramidal neurons of the CA1 region provide the only hippocampal output to cortical targets, this activity must have functional significance. We just have to figure out what it is” (György Buzsáki, 2019). To take a few steps in this direction, our study aimed to zoom in on the HC-DN correspondence in the context of subjective social isolation.

Indeed, recent evidence has shown that DN grey matter morphology is strongly associated with loneliness (Spreng et al., 2020). Analogously, the hippocampus has been discussed to be particularly affected by social isolation in various animal species (Biggio et al., 2019; Ibi et al., 2008; Kogan et al., 2000; Silva-Gomez et al., 2003). Moreover, researchers have been able to perform these invasive studies on the animal hippocampus at a subregion and even single-cell level. This fine-grained resolution is challenging to realize with respect to DN subregions in the living human brain. Since the HC and DN subregions are closely related, studying their structural divergence in lonely individuals provides a window that can offer refined understanding of the association between loneliness and its brain basis. By joint analysis of the HC and DN, the existing in-depth knowledge of the animal hippocampus may help elucidate the nature of potentially human-specific DN subregions. Moreover, in addition to high-resolution structural brain imaging and social isolation information, the availability of genetic data allowed us to also investigate the differing contributions of genetic influences on structural brain patterns related to loneliness.

## Material and Methods

### Population data resource

The UK Biobank is a prospective epidemiology resource that offers extensive behavioral and demographic assessments, medical and cognitive measures, as well as biological samples in a cohort of ∼500,000 participants recruited from across Great Britain (https://www.ukbiobank.ac.uk/). This openly accessible population dataset aims to provide brain-imaging for ∼100,000 individuals planned for completion in 2022. The present study was based on the recent data release from February/March 2020. To improve comparability and reproducibility, our study built on the uniform data preprocessing pipelines designed and carried out by FMRIB, Oxford University, UK (Alfaro-Almagro et al., 2018). Our study involved data from 38,701 participants with brain-imaging measures and expert-curated image-derived phenotypes of grey matter morphology (T1-weighted MRI) from 48% men and 52% women, aged 40-69 years when recruited (mean age 55, standard deviation [SD] 7.5 years). The present analyses were conducted under UK Biobank application number 25163. All participants provided informed consent. Further information on the consent procedure can be found elsewhere (http://biobank.ctsu.ox.ac.uk/crystal/field.cgi?id=200).

### Loneliness target phenotype

Regarding the loneliness target phenotype, we used the yes/no answer from UK Biobank participants to the question ‘Do you often feel lonely?’ (data field 2020), as a subjective indicator of the quality of social interactions. Measures of social relationship quality represent a widely accepted and commonly investigated component of social embeddedness (Cohen and Hoberman, 1983; Hawkley et al., 2005). Loneliness is more commonly viewed as a subjective feeling of being alone, regardless of social encounter frequency (Cacioppo and Cacioppo, 2014). Conceptually similar and highly correlated scales (Cyranowski et al., 2013) are also contained in other standard measurement tools of social embeddedness, such as the Revised UCLA Loneliness Scale (Hawkley et al., 2005) and the Interpersonal Support Evaluation List (Cohen and Hoberman, 1983). In general, a variety of studies showed single-item measures of social isolation traits to be reliable and valid (Atroszko et al., 2015; Dollinger and Malmquist, 2009; Mashek et al., 2007). The demographic differences between lonely and non-lonely individuals in the UK Biobank have been reported elsewhere (Spreng et al., 2020).

### Brain-imaging and preprocessing procedures

Magnetic resonance imaging scanners (3T Siemens Skyra) were matched at several dedicated data collection sites with the same acquisition protocols and standard Siemens 32- channel radiofrequency receiver head coils. To protect the anonymity of the study participants, brain-imaging data were defaced and any sensitive meta-information was removed. Automated processing and quality control pipelines were deployed (Alfaro-Almagro et al., 2018; Miller et al., 2016). To improve homogeneity of the imaging data, noise was removed by means of 190 sensitivity features. This approach allowed for the reliable identification and exclusion of problematic brain scans, such as due to excessive head motion.

The structural MRI data were acquired as high-resolution T1-weighted images of brain anatomy using a 3D MPRAGE sequence at 1 mm isotropic resolution. Preprocessing included gradient distortion correction (GDC), field of view reduction using the Brain Extraction Tool (Smith 2002) and FLIRT (Jenkinson and Smith 2001; Jenkinson et al. 2002), as well as non-linear registration to MNI152 standard space at 1 mm resolution using FNIRT (Andersson et al. 2007). To avoid unnecessary interpolation, all image transformations were estimated, combined and applied by a single interpolation step. Tissue-type segmentation into cerebrospinal fluid (CSF), grey matter (GM) and white matter (WM) was applied using FAST (FMRIB’s Automated Segmentation Tool, (Zhang et al. 2001)) to generate full bias-field-corrected images. SIENAX (Smith et al. 2002), in turn, was used to derive volumetric measures normalized for head sizes.

For the default network, volume extraction was anatomically guided by the Schaefer-Yeo reference atlas (Schaefer et al., 2017). Among the total of 400 parcels, 91 subregion definitions are provided as belonging to the default network among the 7 canonical networks. For the hippocampus, 38 volume measures were extracted using the automatic Freesurfer sub-segmentation (Iglesias et al., 2015). The allocortical volumetric segmentation draws on a probabilistic hippocampus atlas with ultra-high resolution at ∼0.1mm isotropic. In particular, this tool from the Freesurfer 7.0 suite gives special attention to surrounding anatomical structures to refine the hippocampus subregion segmentation in each participant.

As a preliminary data-cleaning step, building on previous UK Biobank research (Schurz et al., 2021; Spreng et al., 2020), inter-individual variation in brain region volumes that could be explained by nuisance variables of no interest were regressed out: body mass index, head size, head motion during task-related brain scans, head motion during task-unrelated brain scans, head position and receiver coil in the scanner (x, y, and z), position of scanner table, as well as the data acquisition site, in addition to age, age^2^, sex, sex*age, and sex*age^2^. The cleaned volumetric measures from the 91 DN subregions in the neocortex and the 38 HC subregions in the allocortex served as the basis for all subsequent analysis steps.

### Structural variation relationships of the fornix and nucleus accumbens

In the context of HC-DN correspondence, the fornix white-matter tract plays a central role for several important reasons. The fornix constitutes the major output pathway of the hippocampus, carrying information unidirectionally towards the neocortical mantle (Aggleton et al., 2015). Our previous research has shown that the fornix is the major white-matter tract whose microstructural variation is by far most predictable based on DN subregion volumes alone (Kernbach et al., 2018). This fiber pathway has also recently been shown to be linked to loneliness (Spreng et al., 2020). Additionally, on its way, the fornix carries axons from hippocampus neurons to the nucleus accumbens (Aggleton et al., 2015; Friedman et al., 2002) – a core node of the reward circuitry, which has been implicated in the sequelae of loneliness, such a substance abuse (Bzdok and Dunbar, 2020).

For these reasons, we have conducted preliminary regression analyses for the fornix (UKB data fields 25095, 25094, and 25061), and for the nucleus accumbens (UKB data fields 25024 and 25023). The ancillary analyses explored their structural relationships with the subregions composing our DN atlas, and those composing our HC atlas (cf. above). The model specification for these regression analyses was as follows:

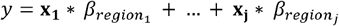

where *β* denote the slope parameters corresponding to the brain volumes **x** for all subregion volumes of either the DN atlas or the HC atlas (z-scored across participants), *y* denotes the mean FA of the fornix or the volume of the nucleus accumbens for the UK Biobank participants.

### Analysis of co-variation between hippocampus subregions and default-network subregions

As the central step of the analytical workflow, we sought dominant regimes of structural correspondence – signatures or “modes” of population co-variation that provide insights into how *structural variation among the segregated HC* can explain *structural variation among the segregated DN*. Canonical correlation analysis (CCA) was a natural choice of method to interrogate such a multivariate inter-relation between two high-dimensional variable sets (Bzdok et al., 2019; Wang et al., 2020; Witten et al., 2009). A first variable set *X* was constructed from the DN subregion volumes (number of participants x 91 DN parcels matrix). A second variable set *Y* was constructed from the HC subregion volumes (number of participants x 38 HC parcels matrix):

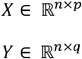

where *n* denotes the number of observations or participants, *p* is the number of DN subregions, and *q* is the number of HC subregions. Each column of the two data matrices was z-scored to zero mean (i.e., centering) and unit variance (i.e., rescaling). CCA addresses the problem of maximizing the linear correlation between low-rank projections from the two variable sets or data matrices. The two sets of linear combinations of the original variables are obtained as follows:

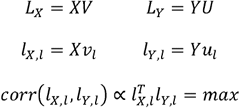

where *V* and *U* denote the respective contributions of *X* and *Y, L_X_* and *L_X_* denote the respective latent ‘modes’ of joint variation based on patterns derived from *X* and patterns derived from Y, *l_X,l_* is the lth column of *L_X_*, and *l_Y,l_* is the lth column of *L_Y_*. The goal of our CCA application was to find pairs of latent vectors *l_X,l_* and *l_Y,l_* with maximal correlation in the derived latent embedding. In an iterative process, the data matrices *X* and *Y* were decomposed into *L* components, where *L* denotes the number of modes given the model specification. In other words, CCA involves finding the canonical vectors *u* and *v* that maximize the (symmetric) relationship between a linear combination of DN volume expressions *(X)* and a linear combination of HC volume expressions *(Y)*. CCA thus identifies the two concomitant projections X*v_l_* and Y*u_l_*. These yielded the optimized co-occurrence between patterns of subregion variation inside the segregated DN and patterns of subregion variation inside the segregated HC across participants.

In other words, each estimated cross-correlation signature identified a constellation of within-DN volumetric variation and a constellation of within-HC volumetric variation that go hand-in-hand with each other. The set of k orthogonal modes of population co-variation are mutually uncorrelated by construction (Wang et al., 2020). They are also naturally ordered from the most important to the least important HC-DN co-variation mode based on the amount of variance explained between the allocortical and neocortical variable sets. The first and strongest mode therefore explained the largest fraction of joint variation between combinations of HC subregions and combinations of DN subregions. Each ensuing cross-correlation signature captured a fraction of structural variation that is not explained by one of the *k* − 1 other modes. The variable sets were entered into CCA after a confound-removal procedure based on previous UK Biobank research (cf. above).

### Group difference analysis

For the derived population modes of HC-DN co-variation, we then performed a rigorous group contrast analysis in the context of loneliness. We aimed to identify which anatomical subregions show statistically defensible deviation in the lonely group compared to the control group. We carried out a principled test for whether the solution vector obtained from the CCA (i.e., canonical vectors, cf. above) in the lonely group is systematically different from the solution vector in the control group.

More specifically, following previous UK Biobank research (Schurz et al., 2021), we carried out a bootstrap difference test of the CCA solution from the lonely vs. non-lonely group (Efron and Tibshirani, 1994). In 100 bootstrap iterations, we randomly pulled participant samples with replacement to build an alternative dataset (with the same sample size) that we could have gotten. We subsequently performed CCA in parallel by fitting one separate model to each of the two groups. In each resampling iteration this approach thus carried out a separate estimation of the doubly multivariate correspondence between HC subregions and DN subregions in each particular group. The two distinct CCA solutions from each iteration were then matched mode-by-mode regarding sign invariance and mode order. Canonical vectors of a given mode that carried opposite signs were aligned by multiplying one with -1. The order of the CCA modes was aligned based on pairwise Pearson’s correlation coefficient between the canonical vectors from each estimated CCA model. After mode matching, we directly estimated the resample-to-resample effects by elementwise subtraction of the corresponding canonical vectors of a given mode *k* between the two groups. We finally recorded these difference estimates for each vector entry (each corresponding to the degree of deviation in one particular anatomical subregion). The subregion-wise differences were ultimately aggregated across the 100 bootstrap datasets to obtain a non-parametric distribution of group contrast estimates.

We thus propagated the variability attributable to participant sampling into the computed uncertainty estimates of group differences in the UK Biobank population cohort. Statistically relevant alteration of anatomical subregions in loneliness were determined by whether the two-sided confidence interval included zero or not according to the 10/90% bootstrap-derived distribution of difference estimates (Schurz et al., 2021). In an approach that is faithful to our multivariate analytical strategy and research question. This non-parametric approach directly quantified the statistical uncertainty of how loneliness is manifested in specific subregions of the HC-DN axis.

### Analysis of how individual expressions of hippocampus-DN co-variation are linked to the genetic predisposition for loneliness

Polygenic risk scores (PRS) is a genome-wide analysis technique that has been shown to successfully quantify individuals genetic predisposition for a variety of phenotypes. The approach has become especially potent for complex phenotypes that implicate tens of thousands of single nucleotide polymorphisms (SNPs) with individually small effect sizes, such as major psychiatric diseases (Elliott et al., 2020; HLA-C, 2009; Inouye et al., 2018; Khera et al., 2018; Kuchenbaecker et al., 2017). PRS have also come to be a sharp tool for heritability analyses due to the advent of large population datasets, such as the UK Biobank (Choi et al., 2020; Wray et al., 2021). Such data resources have allowed the investigation of the relationship between SNP variation and inter-individual differences in a particular phenotype, which includes neuroimaging-derived phenotypes (Elliott et al., 2018; Smith et al., 2021). For the purpose of the present study, we have constructed PRS models for the loneliness trait. The subject-specific risk scores were then regressed onto our expressions of HC-DN modes (i.e., canonical variates). The integrative imaging-genetics approach allowed disentangling which mode expressions showed reliable relationships to the heritability of loneliness as attributable to thousands of genetic variants.

As is common for PRS analysis workflows, summary statistics from previously conducted genome-wide association studies (GWAS) on our target phenotypes were used as the backdrop to determine how several hundred thousand SNPs are associated with the loneliness trait. The summary statistics for loneliness were obtained from a GWAS that was conducted as part of the Psychiatric Genomics Consortium. Quality control was implemented by excluding SNPs with a minor allele frequency of less than 1%, as well as excluding SNPs with imputation information score of less than 0.8. Mismatching, duplicate, and ambiguous SNPs were also disregarded from further analysis. Quality control on the base data also involved excluding individuals with a difference between reported sex and that indicated by their sex chromosomes, and removing overlapping samples.

The quality-controlled summary statistics were used as starting point for the PRS model that was built and applied using the PRSice framework (http://www.prsice.info). This software tool uses the available collection of effect sizes of candidate SNPs to form single-subject predictions of genetic predisposition for a phenotype of interest. In particular, this tool determined the optimal PRS model based on the UK Biobank participants (training data, n=253,295) of European ancestry who did not provide any brain-imaging data (at the time of study). This model training step involved automated adjustments, such as identifying ideal clumping and pruning choices, to select the thresholds that decide which SNPs are included in the PRS model. Subsequently, once optimized, the final PRS model was then used to predict the genetic predisposition for each of 23,423 UK Biobank participants of European ancestry *with* brain-imaging data (test data). This PRS model consisted of the additive effects of weighted SNPs, whereby the weighted sum of the participants’ genotypes was computed as follows:

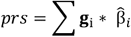

where g_i_ denotes an individual’s genotype at SNP *i* (values 0, 1, or 2), 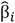 is the obtained point estimates of the per-allele effect sizes at SNP *i*, and *j* is a particular individual (Choi et al., 2020).

Finally, Bayesian linear regression was used to regress the subject-specific predictions of genetic liability for the loneliness trait onto the participant expressions of HC-DN co-variation modes. More specifically, the individuals in the top 5% predictions (i.e., highest PRS estimates) and the individuals in the bottom 5% predictions (i.e., lowest PRS estimates) were considered in a Bayesian logistic regression model with mode expressions serving as input variables (Fan et al., 2019; Lecarpentier et al., 2017; Meisner et al., 2020). In this multiple regression setup, PRS was regressed against each of the 25 canonical variates (linearly uncorrelated by construction) for each individual on the hippocampus side. An analogous multiple regression model was estimated for the (uncorrelated) 25 canonical variates from the DN side. The fully Bayesian model specification for these regression analyses was as follows:

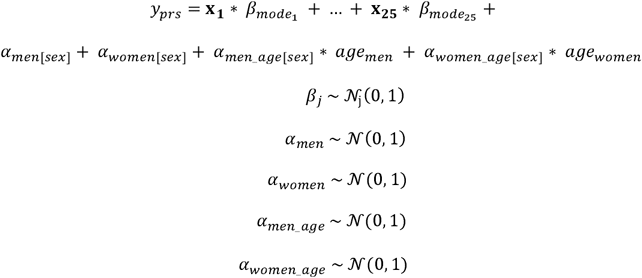

where β_*j*_ denote the slopes for the subject-specific 25 mode expressions as canonical variates x_j_, *y_prs_* denotes the PRS estimates of each participant. Potential confounding influences were acknowledged by the nuisance variables α, which accounted for variation that could be explained by sex and (z-scored) age. O nce the Bayesian model solution was approximated using Markov chain Monte Carlo sampling, it yielded fully probabilistically specified posterior parameter distributions for each β coefficient corresponding to one of the signatures of allocortical-neocortical co-variation (cf. above). The association with trait heritability of a mode expression was then determined based on how robustly their corresponding model coefficients deviated from 0 (e.g., >95% of model coefficient posterior probability excluded a value of 0).

## Results

### Rationale

Previous brain-imaging studies aimed at brain parcellation have typically investigated either the default network alone (e.g., Schurz et al., 2014) or the hippocampus alone (e.g., Plachti et al., 2019). Additionally, there is still insufficient work relating animal studies on anatomically defined subregions of the hippocampus to what can be reliably measured in the human medial temporal lobe using brain-imaging techniques. Advances in the automatic segmentation of the hippocampus using ex vivo brain-imaging (Iglesias et al., 2015; Wisse et al., 2017) now allow the reliable assessments of microanatomically defined hippocampus subregions in a way that scales to the ∼40,000 UK Biobank Imaging cohort. This enables deeper analyses of the principled inter-relationships between the evolutionarily more conserved hippocampus of the allocortex and the default network of the recently expanded neocortex. By leveraging a framework for high-dimensional co-decomposition at a fine-grained subregion resolution, we here investigate the structural deviations of the HC-DN co-variations signatures that characterize loneliness.

### Structural variation relationships of the fornix

In a preliminary set of exploratory analyses, we sought to elucidate whether individual subregion volumes of the hippocampus could distinctly explain microstructural variation of the fornix white-matter tract ⍰ an important fiber pathway known to carry axons from the allocortical hippocampus to the DN. To this end, we performed a multiple regression analysis to regress the structural integrity of the fornix (mean FA, diffusion MRI) onto 38 hippocampal subregion volumes. We found that the bilateral CA1 body dominates in explaining variation in fornix integrity, followed by the molecular layer (ML) head of the left hippocampus (Figure 1A). Smaller positive effects were found in the bilateral subiculum body, bilateral presubiculum body, bilateral fimbria, right ML head, and right CA2/3. We found negative effects in the bilateral hippocampal fissure, left parasubiculum, bilateral CA4 head, as well as bilateral CA4 body. These findings indicate that several hippocampal subregions have unique structural relationships with fornix integrity. In a variant of this analysis, rather than marginal or partial association strengths, we computed pairwise Pearson correlation strengths between fornix integrity and each hippocampal subregion. The hippocampus subregions with the largest correlation coefficients rho were also the subregions with the greatest amount of explained variance in the multiple regression analysis (Supplementary Figure 1, https://doi.org/10.6084/m9.figshare.15060684). Thus, from convergent evidence across two different analyses of how hippocampus subregions track fornix variation, the CA1 body showed the strongest positive association with the fornix, while the hippocampal fissure displayed the strongest negative association.

**Figure 1.**
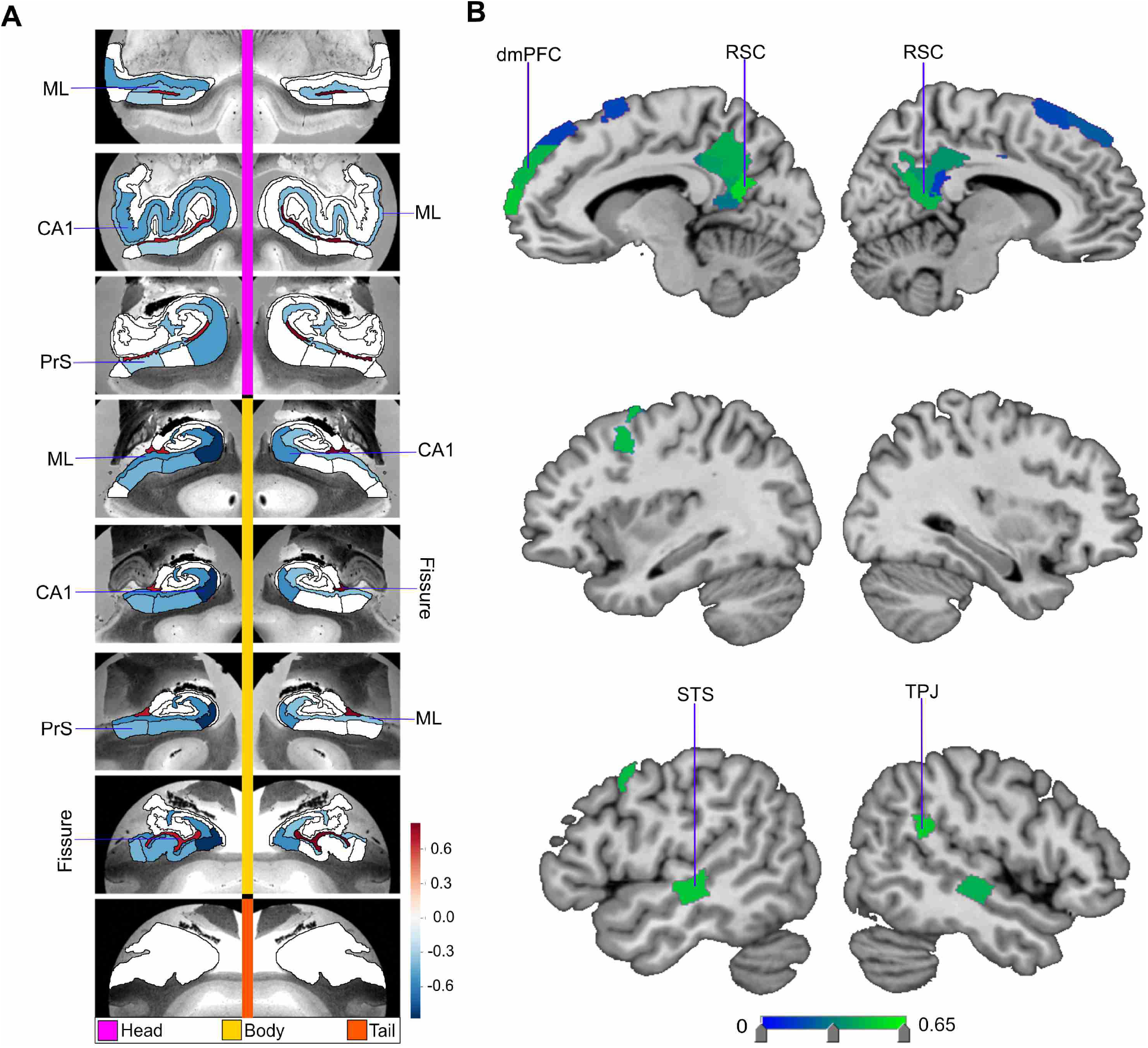
Structural correlates of the fornix white-matter tract. How can microstructural variation of the fornix be explained as a function of subregion volume variations in the hippocampus (left) or the default network (right)? To address these two questions, we have computed separate exploratory multiple regression analyses based on 38,701 UK Biobank participants. The regression parameter weights corresponding to each specific subregion (hot and cold colors) outline the structural associations with fornix integrity exclusively explained by that particular subregion. **A** shows the hippocampus-fornix model based on 38 subregions from the hippocampus segmentation (Iglesias et al., 2015). The parameter weights indicate the variation explained in the fornix specifically by each hippocampal subregion, mapped onto 8 consecutive coronal sections of the left and right hippocampus from anterior (top) to posterior (bottom) direction (hot/cold colors = positive/negative volume association). The subregions with the strongest positive volume effect are CA1 body of the left and right hippocampus, as well as left molecular layer head. The strongest negative relations with fornix microstructure are found in left and right hippocampal fissures. **B** shows the default network-fornix model based on 91 subregions in the Schaefer-Yeo parcellation of the default network. The parameter weights indicate the fornix variation explained specifically by each default network subregion. The subregions with the strongest marginal volume effect amongst DN subregions are bilateral posterior cingulate cortex (PCC), bilateral retrosplenial cortex (RSC), and left dorsal-medial prefrontal cortex (dmPFC). Overall, these results suggest that fornix architecture has differentiated structural relationships with subregions of the hippocampus and midline DN. PrS = presubiculum, Sub = subiculum, DG = GC-DG-ML, ML = molecular layer; vmPFC = ventromedial prefrontal cortex, OFC = orbitofrontal cortex.

We next sought to regress variation of fornix integrity onto 91 subregion volumes of the DN (Figure 1B). The subregions with the strongest positive effects included the left dorsal mPFC (dmPFC) and the right retrosplenial cortex (RSC). There was also a small effect in the left RSC, bilateral ventromedial prefrontal cortex (vmPFC) and left dorsolateral prefrontal cortex (dlPFC). Conversely, we located the strongest negative effects to the bilateral posterior cingulate cortex (PCC). Further, in the companion analysis using Pearson correlation between each DN subregion and the fornix, mPFC subregions had in general strong positive effects (Supplementary Figure 2, https://doi.org/10.6084/m9.figshare.15060684). On the other hand, PCC/RSC subregions had weak or negative effects (Supplementary Figure 2, https://doi.org/10.6084/m9.figshare.15060684). These findings thus made apparent the subregion-specific relationships of how DN subregions track fornix integrity in our UKB cohort: with lateral DN subregions showing more subtle effects, and midline DN subregions showing the most prominent effects.

### Structural variation relationships of the nucleus accumbens

Using the identical analysis approach (cf. above), we next explored how the volume of the nucleus accumbens (NAc) can be explained as a function of hippocampal subregion volumes. These multiple regression analyses revealed that bilateral CA1 body and right molecular layer head subregions explained the most variation in NAc volume (Supplementary Figure 2A, https://doi.org/10.6084/m9.figshare.15060684). Small positive effects were also found in CA2/3 head of the right hippocampus, left molecular layer head, and bilateral hippocampus amygdala transition area (HATA). Conversely, there was a strong negative association of the NAc with bilateral hippocampal fissure, as well as with left parasubiculum. Interrogating the HC- NAc relationships thus showed distinct explanatory effects amongst subregions. In particular, the hippocampal subregions which explained the most variation in NAc recapitulated those which also best explained fornix integrity (cf. above). To further investigate the associations of the NAc, we next computed gross pairwise Pearson correlation coefficients of each hippocampal subregion with the NAc. We observed a noticeably stronger correlation for CA1 head than any other subregion (Supplementary Figure 3, https://doi.org/10.6084/m9.figshare.15060684); although CA1 head showed a negligible effect in the multiple regression analysis of the NAc on hippocampus subregions. Thus, while CA1 head on its own has a strong correlation with the NAc, when considered in a joint model with all other hippocampal subregions in our atlas, it explained little NAc variation in a unique way.

This observation suggests that distinct hippocampal subregions, other than CA1, make separate contributions that together better explain volume variation in the NAc.

To complement these hippocampus analyses, the relationship between the NAc and DN subregions was also investigated with a dedicated multiple regression analysis. The subregions with the largest explained variance in NAc volume included right orbitofrontal cortex (OFC), left OFC, left posterior superior temporal sulcus (pSTS), left ventrolateral prefrontal cortex (vlPFC), and left vmPFC (Figure 2B). Conversely, strong negative associations with NAc volume were located to the left and right vmPFC, left and right PCC, as well as right temporal pole. In a Pearson correlation analysis of each DN subregion with the NAc, we also observed that mPFC subregions had generally strong correlations with the NAc, and PCC/RSC subregions had generally weak correlations (Supplementary Figure 4, https://doi.org/10.6084/m9.figshare.15060684). Overall, in contrast to our HC-centric analyses for the fornix and NAc (cf. above), DN subregions were found to have individually varying relationships with the NAc which noticeably drew a different picture than our results obtained for the fornix-DN relationship.

**Figure 2.**
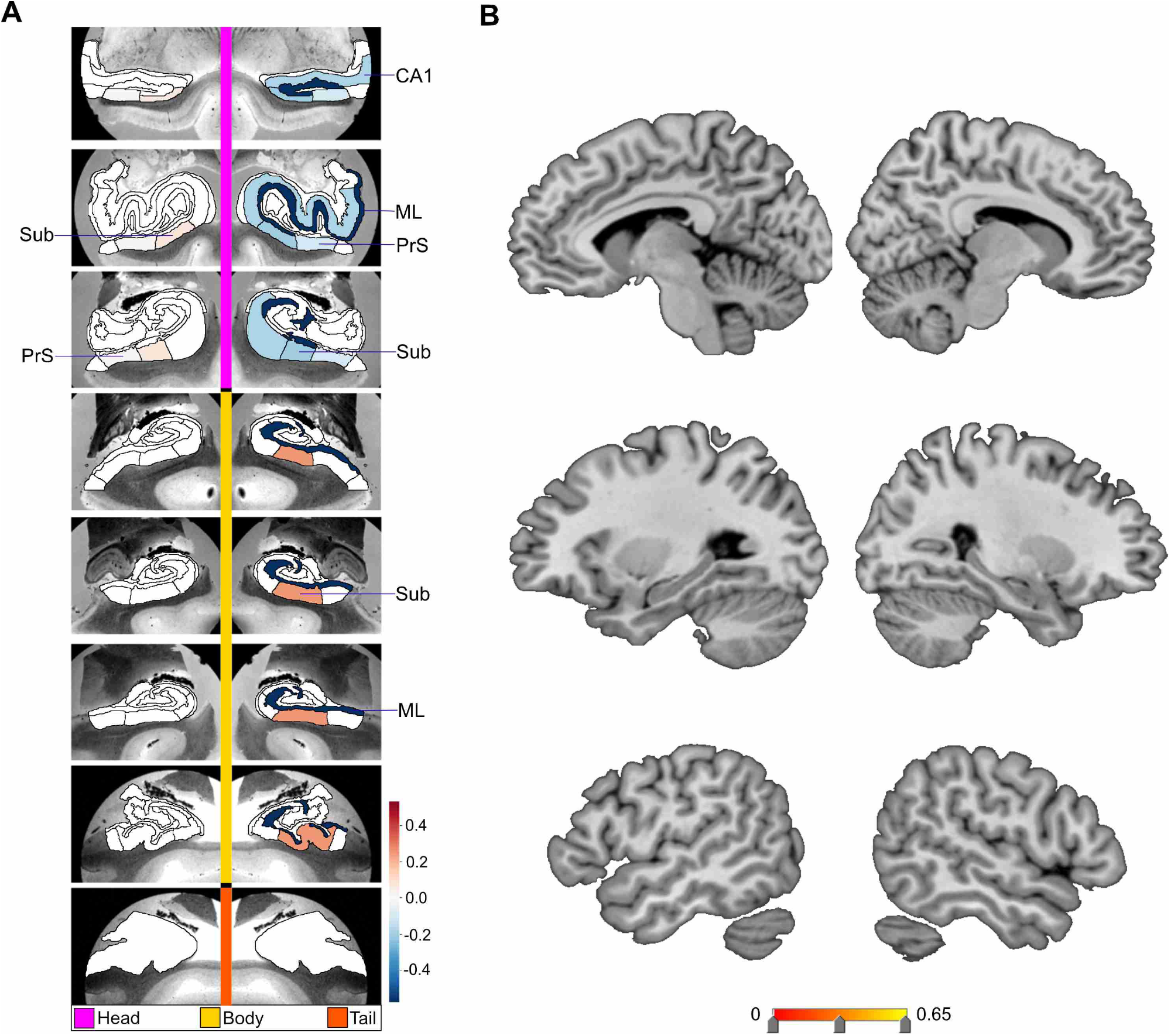
Structural correlates of the nucleus accumbens. Regression parameter weights of various subregions in separate exploratory multiple regression analyses outline the volume effects on the nucleus accumbens (NAc), due exclusively to each subregion in either the hippocampus or default network. **A** shows the hippocampus-NAc model mapped onto 8 consecutive coronal slices of the left and right HC in the anterior (top) to posterior (bottom) direction. The parameter weights indicate the structural variation uniquely explained in the nucleus accumbens specifically by each hippocampal subregion (hot/cold colors = positive/negative volume association). The subregions with the strongest positive volume effect are CA1 body of the left and right hippocampus, as well as right molecular layer head ⍰ in line with our analogous analyses on the fornix (Figure 1A). The strongest negative volume effects are left and right hippocampal fissures. **B** shows the default network-NAc model. The parameter weights indicate the variation explained in the NAc specifically by each default network subregion (warm/cold colors = positive/negative volume association). The subregions with the strongest volume effect amongst default network subregions are bilateral orbitofrontal cortex (OFC), left posterior superior temporal sulcus (pSTS), and left ventro-lateral prefrontal cortex (vlPFC). Overall, these results suggest there are diverse structural relationships between the NAc and hippocampal subregions, which are substantially similar to the structural relationships of the fornix (Figure 1). Yet, the relationship of the NAc with default network subregions shows substantially different volume effect sizes for the most prominent cortical regions when compared to the fornix. PrS = presubiculum, HATA = hippocampal amygdala transition area, DG = GC-DG-ML, ML = molecular layer; vmPFC = ventromedial prefrontal cortex.

### Structural co-variation between hippocampus and default network at the subregion level

In our core analysis, we explored the principled signatures of structural co-variation between the full set of 38 hippocampal subregions and the full set of 91 DN subregions. The concurrent patterns of subregion variation within the hippocampus and within the DN were computed using doubly multivariate pattern-learning analysis. In so doing, we achieved a co- decomposition of hippocampal subregion volumes and DN subregion volumes. Each of the top 25 modes of co-variation was characterized based on how much of joint variance a particular signature explained: with the most explanatory signature (mode 1) achieving a canonical correlation of rho = 0.51 (measured as Pearson’s correlation coefficient) (Supplementary Table 1, https://doi.org/10.6084/m9.figshare.15060684). The second most explanatory signature (mode 2) achieved a canonical correlation of rho = 0.42, the third rho = 0.39, the fourth rho = 0.31, the fifth rho = 0.27, and the sixth rho = 0.23; through to the twenty-fifth signature which had rho = 0.06 (Supplementary Table 1 for full list, https://doi.org/10.6084/m9.figshare.15060684). This analysis thus established the scaffold for all subsequent analyses that delineates how multiple complementary hippocampal patterns co- vary hand-in-hand with DN patterns.

### Differences in the hippocampus-default network co-variation in loneliness

Based on the identified population signatures of HC-DN co-variation, we investigated the neurobiological manifestations of loneliness in our UKB sample. This was accomplished by examining robust subregion-level divergences in how hippocampal patterns are co-expressed with DN patterns between groups of lonely vs. non-lonely participants. To this end, we first analyzed loneliness by a rigorous group difference analysis between the structural patterns of co-variation in the lonely and non-lonely groups. This revealed the precise subregions contributing to the structural HC-DN co-variation that systematically diverged between the two groups, for each mode of the CCA.

We uncovered a multitude of modes with systematic group differences in either specific HC and/or DN subregions. We also found modes with no significant structural divergences in any hippocampal or DN subregion. From here on, a subregion which was observed to have a robustly different weighting within a modes canonical vector, between the lonely and non-lonely groups, is termed a ‘hit’ (i.e., an observed structural divergence in lonely individuals). Across all 25 examined modes, the group contrasts amounted to 28 total hits for HC subregions and 40 total hits for DN regions. Most of these subregion hits occurred in earlier modes. For example, in the first mode we found 26 DN hits (60% of the DN total), and in the first three modes we found 22 HC subregion hits (78.6% of the HC total). We also noted specific subregions with repeated hits across modes: The largest number of hits was in molecular layer body (4 hits), molecular layer head (3), CA1 head (3), CA1 body (3), and presubiculum head (3). In the DN, the greatest number of hits was observed in midline subregions (77.5% of the DN total), such as the RSC and mPFC. We observed few hits in lateral temporal or parietal subregions, with a total number of hits in 5 temporal (12.5%), 13 PFC (32.5%), 2 parietal (5%), and 20 PCC/RSC (50%) subregions.

The constellation of structural divergences between lonely and non-lonely groups in mode 1 (Figure 3) provided a rough portrait of the hits that typically emerged across the next 24 modes. In mode 1 we observed hits in bilateral CA1 head and left CA1 body, the adjacent left presubiculum head and body, as well as the bilateral subregions internal to the hippocampus (ML and fissure). Concurrently, the DN hits for mode 1 were clustered in adjacent subregions in the bilateral PCC/RSC region and mPFC (2 temporal, 8 PFC, 1 parietal, 15 PCC). Additionally, the subregions with hits that played an especially strong role in the dominant mode’s patterns were bilateral CA1 body, bilateral fissure, bilateral RSC, and left dmPFC. In contrast, for the second mode of the HC-DN co-variation, we observed hits only in the left parasubiculum and left ML head, with no hits in DN regions (Supplementary Figure 5, https://doi.org/10.6084/m9.figshare.15060684). For mode 3, we observed 8 HC hits and no DN hits (Figure 4); for mode 4, 1 hit in the HC tail and no DN hits (Supplementary Figure 6, https://doi.org/10.6084/m9.figshare.15060684); mode 5, 1 HC and 1 DN hit (Figure 5); mode 6, no hits; mode 7, no HC hits and 1 hit in the right tempo-parietal junction (Supplementary Figure 7, https://doi.org/10.6084/m9.figshare.15060684); mode 8, HC hits in 3 subregions ⍰ the left CA1 head, left CA2/3 head, and left GC-ML-DG head, and no DN hits (Supplementary Figure 8, https://doi.org/10.6084/m9.figshare.15060684); mode 9, 1 HC hit and 7 DN hits (Figure 6). We observed no hits in modes 10 through 13. In mode 14 we found no HC hits and 3 DN hits: left superior temporal sulcus (STS), right anterior cingulate cortex (ACC), and left RSC (Supplementary Figure 8, https://doi.org/10.6084/m9.figshare.15060684). In mode 15, we found no HC hits and 2 hits in DN subregions: bilateral PCC (Supplementary Figure 9, https://doi.org/10.6084/m9.figshare.15060684). Of note, we observed no hits in any of the modes between 16-25. Thus the dominant modes, which account for more population covariance, are most strongly coupled with loneliness.

**Figure 3.**
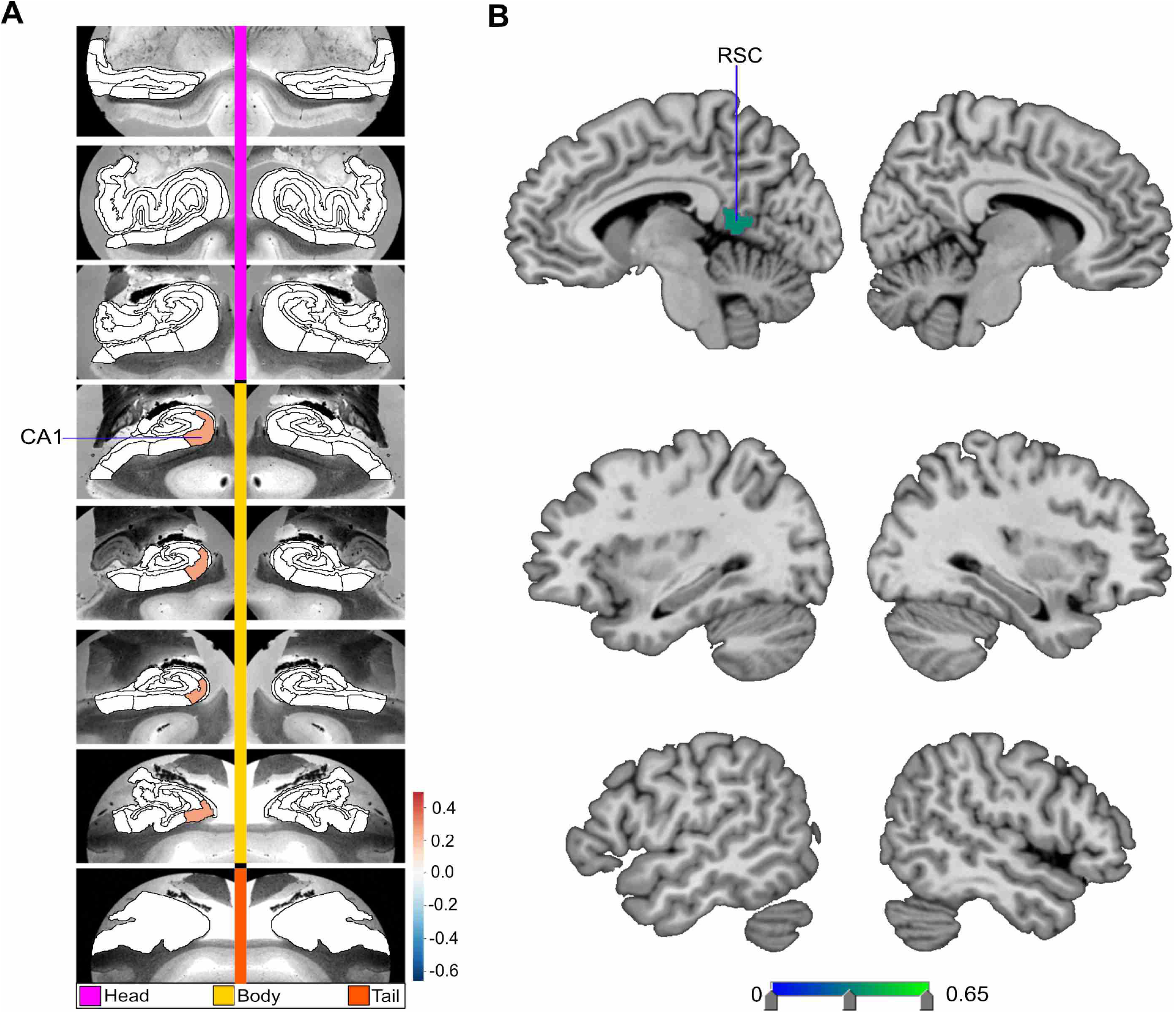
Loneliness is associated with distinct divergences in hippocampus-default network co-variation. We explored the structural co-variation between the 38 subregions of the hippocampus (HC) and 91 subregions of the default network (DN), by means of a co- decomposition based on a canonical correlation analysis (CCA). We subsequently determined how the ensuing subregion patterns diverged in individuals with loneliness. Shown here are the subregion divergences in mode 1 of hippocampus-DN covariation. Mode 1 of the CCA solution acheives the most explanatory co-variation, with a canonical correlation of rho = 0.51. **A** shows the HC subregion patterns (left, one canonical vector of mode 1) with parameter weights that robustly diverge between lonely and non-lonely groups in mode 1; mapped onto 8 consecutive coronal slices of the left and right HC in the anterior (top) to posterior (bottom) direction. **B** shows the DN subregions patterns (right, other canonical vector of mode 1) that robustly diverged between the lonely and non-lonely groups. Overall, within the dominant structural co- variation pattern between the HC and the DN there are specific subregions whose volumes systematically diverge in lonely individuals. The most pronounced structural divergences are in bilateral CA1 body and hippocampal fissure, as well as sub-regions bilaterally in the posterior cingulate cortex (PCC), retrosplenial cortex (RSC), and dorsomedial prefrontal cortex (dmPFC). Thus, specific HC and DN anatomical subregions are preferentially linked loneliness. STS = superior temporal sulcus, TPJ = temporoparietal junction, ML = molecular layer, PrS = presubiculum, Sub = subiculum.

**Figure 4.**
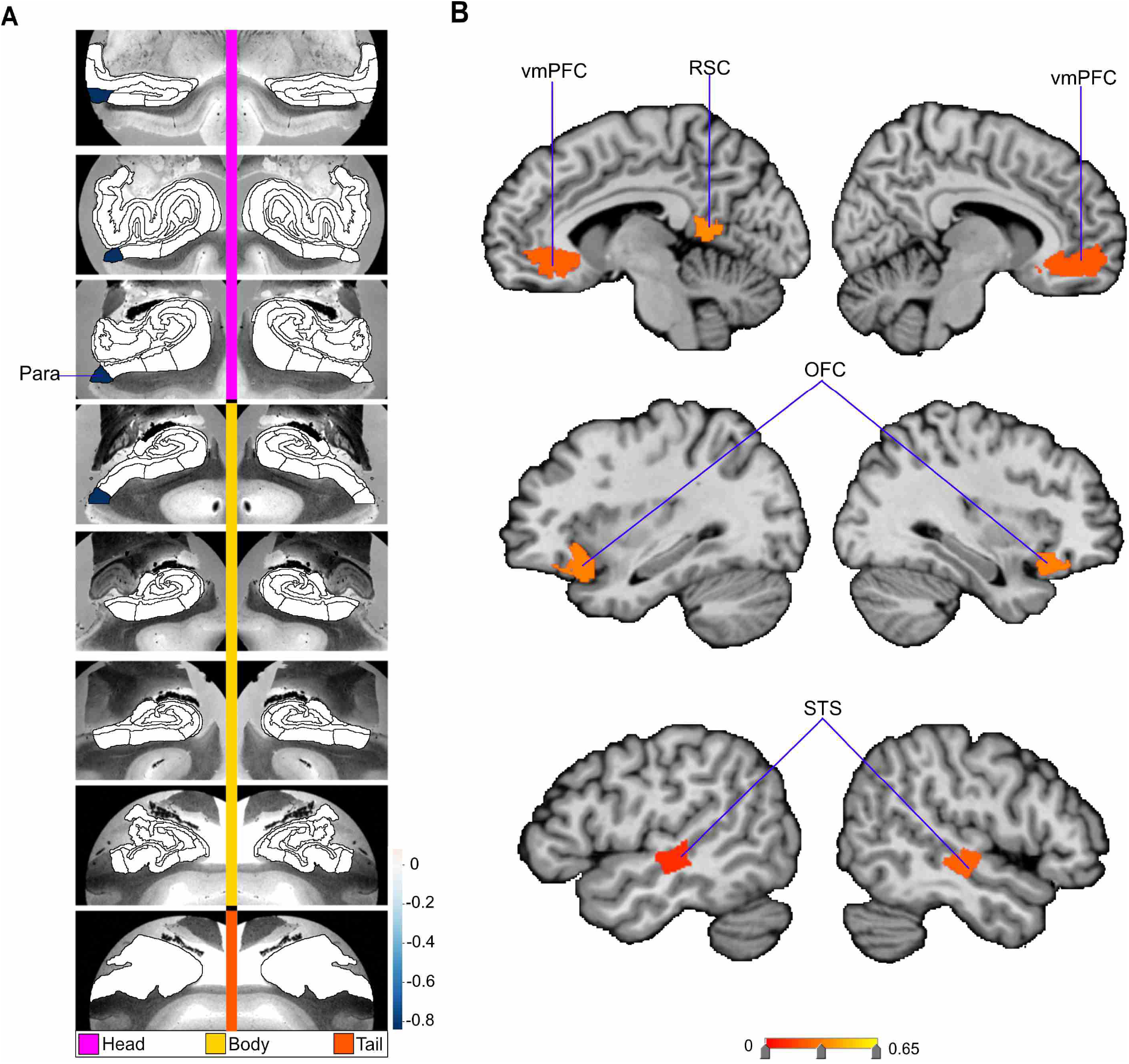
Divergences in CA1, subiculum, presubiculum and molecular layer subregion volumes are preferentially associated with loneliness. Shown here are the subregion divergences in mode 3 of hippocampus-DN co-variation. Mode 3 of the CCA solution acheives the third most explanatory hippocampal-default network co-variation, with a canonical correlation of rho = 0.39. **A** exhibits the hippocampal (HC) subregion patterns (left, one cannical vector of mode 3) with paramater weights that robustly diverge between lonely and non-lonely groups in mode 3. **B** shows the default network (DN) subregion patterns (right, other canonical vector of mode 3) of the default network (right) that robustly diverge between lonely and non- lonely groups. The most pronounced divergences in the loneliness group are in right molecular layer head and body, right subiculum head and body, as well as right CA1 head. Thus, similar to mode 1, within the third most explanatory pattern of HC-DN co-variation, loneliness is concomitant with hippocampal divergences in CA1, subiculum (Sub), presubiculum (PrS), and moleuclar layer (ML).

**Figure 5.**
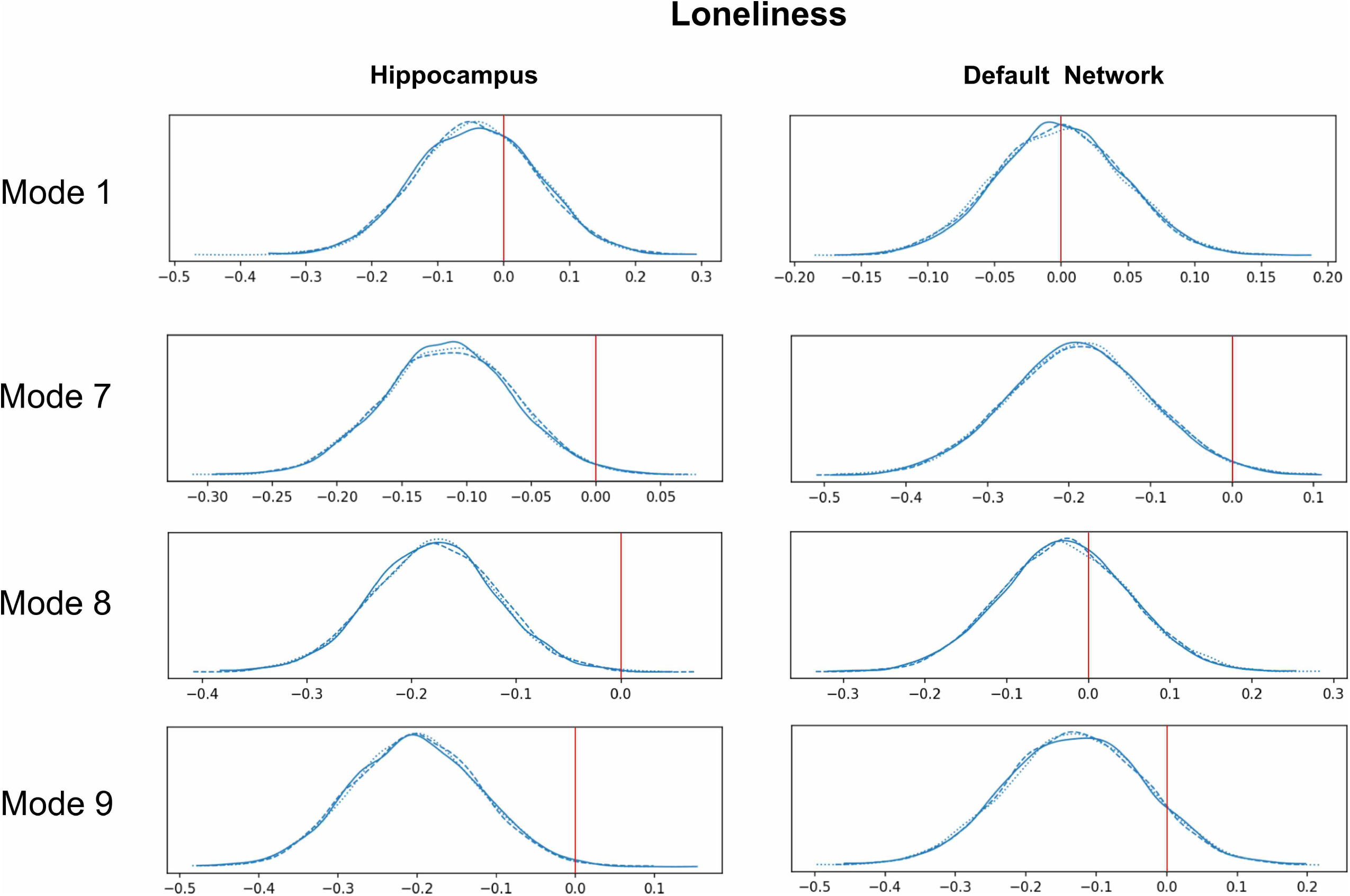
Loneliness is associated with divergences in left hemisphere CA1 and retrosplenial cortex. Shown here are the subregion divergences in mode 5 of hippocampus-DN co-variation. Mode 5 of the CCA solution acheives the third most explanatory hippocampal-default network co-variation, with a canonical correlation of rho = 0.27. **A** exhibits the hippocampal (HC) subregion patterns (left, one cannical vector of mode 5) with paramater weights that robustly diverge between lonely and non-lonely groups in mode 5. **B** shows the default network (DN) subregion patterns (right, other canonical vector of mode 5) of the DN (right) that robustly diverge between lonely and non-lonely groups. The only subregion divergences associated with loneliness are in left CA1 body, and the left retrosplenial cortex (RSC). These results accentuate the selectivity of the subregion divergences within a particular mode, and highlights CA1 and RSC in loneliness.

**Figure 6.**
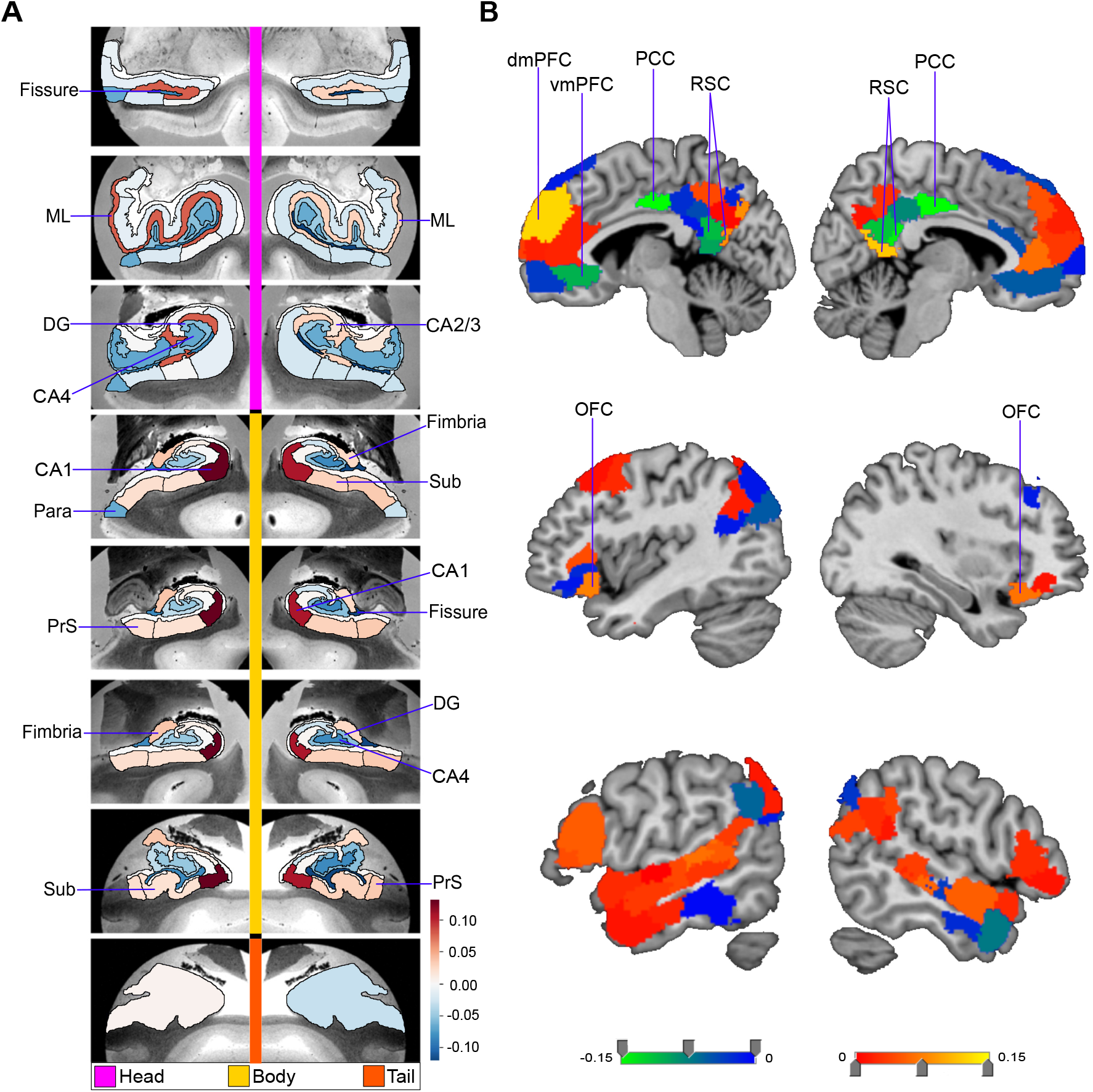
Loneliness is associated with subregions divergences in parasubiculum and a distributed set of bilateral default-network subregions. Shown here are the subregion divergences in mode 9 of hippocampus-DN covariation. Mode 9 of the CCA solution acheives the third most explanatory hippocampal-default network co-variation, with a canonical correlation of rho = 0.18. **A** exhibits the hippocampal (HC) subregion patterns (left, one cannical vector of mode 9) with paramater weights that robustly diverge between lonely and non-lonely groups in mode 9. **B** shows the default network (DN) subregion patterns (right, other canonical vector of mode 9) of the DN (right) that robustly diverged between lonely and non-lonely groups. Overall, there are divergences for lonely indviduals in left parasubiculum (Para), as well as bilateral ventromedial prefrontal cortex (vmPFC), left retrosplenial cortex (RSC), bilateral orbitofrontal cortex (OFC), and bilateral superior temporal sulcus (STS). These results emphasize the bilateral tendency of the DN divergences observed for loneliness. The results also highlight a coherent set of reward related subregions ⍰ as most of the subregions identified are also found to have a strong volummetric relationship with the nucleus accumbens (Figure 2).

These results suggest that specific subregions that play a role in the HC-DN correspondence exhibit systematic diverges in individuals with loneliness. Additionally, our findings specify within which context of structural dependence (mode of co-variation) a subregion diverged between the two groups. Overall, across loneliness’ relationships to our discovered population modes of joint volume variation, CA1 head and body, molecular layer head and body, and presubiculum head showed the largest number of hits for hippocampal subregions. On the other hand, there was a preponderance of hits located to midline cortical structures ⍰ such as the mPFC and PCC/RSC ⍰ for DN subregions. Thus, our population-level findings made apparent that within structural co-variation modes, loneliness was mostly concomitant with divergences in DN midline subregions that were grounded in parallel divergences in the CA1 and ML subregions of our hippocampus atlas.

### Genetic predisposition for loneliness

We finally sought to interrogate whether the uncovered HC-DN co-variation expressions featured systemic relationships with the participants’ liability for loneliness (cf. methods). For this purpose, we first computed polygenic risk score predictions of loneliness risk for our UK Biobank participants. We observed that there was a statistically relevant relationship between loneliness PRS score and participant expressions (i.e., canonical variates) of modes 7, 8, 9 and 22 in the hippocampus, and the participant expressions of mode 7 in the DN (Figure 7). On the whole, there was genetic predisposition linked to the expressions of later modes compared to the earlier modes achieving greater explained variance. This analysis suggested that the expression of the identified HC-DN signatures tracks the purely heritable components of loneliness due to single nucleotide polymorphism. As a matter of course, the signatures not identified likely have contributions that are not due to genetic factors.

**Figure 7.**
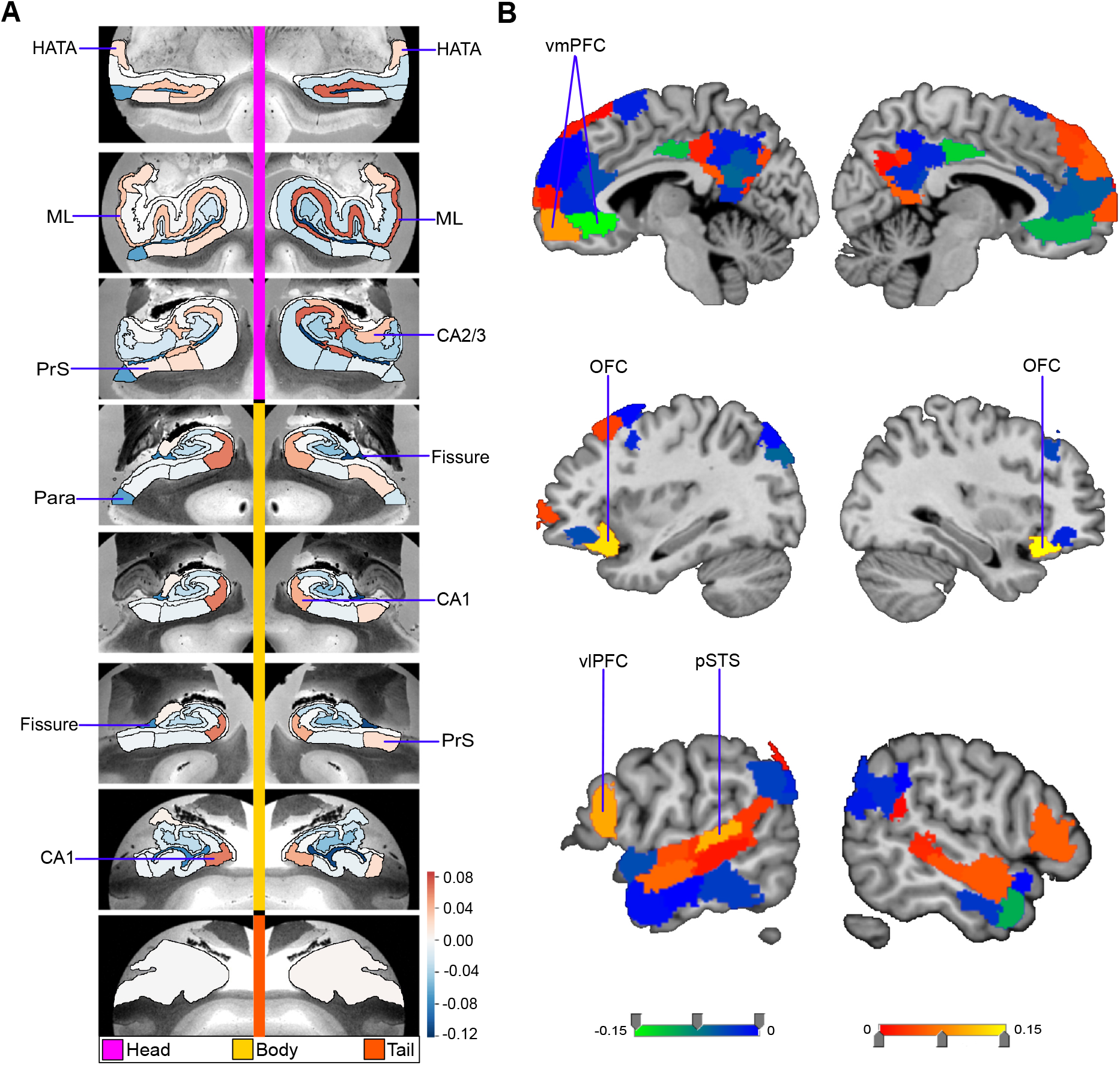
The genetic predisposition for loneliness is associated with the expression of specific hippocampus-default network co-variation patterns. A polygenic risk score (PRS) analysis was conducted to estimate the subject-specific heritable tendency for loneliness based on genome-wide effects in single nucleotide polymorphisms. The subject-specific PRS estimates were then regressed against the expressions of each of the modes of structural co-variation between the hippocampus (HC) and default network (DN). The relevance of the heritability effects was judged based on the posterior parameter distributions inferred by the Bayesian logistic regression model (histograms). The x axis of each plot represents the β coefficient of the model parameter, and the y axis of each plot represents the plausibility of each coefficient value. A value of 0 indicates no association between PRS and inter-individual mode expression. Overall, loneliness predisposition was related to the expression of modes 7 and 9 on the HC side, and to mode 7 on the DN network side. These results demonstrate that specific mode expressions are robustly linked to the predisposition of becoming lonely. The three posterior distribution histograms show convergence across three different Markov chain Monte Carlo runs. Overall, this pinpoints the specific neurobiological signatures in the hippocampus and default network which relate to the heritable components of loneliness.

## Discussion

We have tailored an analytical framework to examine how the structural substrates of the HC-DN correspondence systematically deviate in loneliness. We work towards this goal by directly estimating principled co-variation signatures that delineate how hippocampus subregion volumes are co-expressed with DN subregion volumes in ∼40,000 participants from the UK Biobank cohort. In so doing, our study aimed to refine the understanding of the inter- relationship of the DN with loneliness. By anchoring our analysis in HC-DN co-variation, we aimed to facilitate the interpretation of potentially human-specific DN brain regions based on their hippocampal subregion affiliates. Indeed, the evolutionarily conserved hippocampus has been extensively studied in animal species at a single-cell resolution. This approach can detect patterns that are not discoverable by any analysis of the HC or the DN alone.

The DN has recently been found to be the major brain network that is most closely associated with loneliness (Spreng et al., 2020). However, we found a high degree of heterogeneity in the relationship of individual DN subregions with loneliness in the context of population modes of HC-DN volume co-variation. In particular, we found that especially midline structures of the DN tended to diverge in lonely individuals. Analogously, the hippocampal subregions with the most divergences across modes were CA1 and molecular layer. The allocortical-neocortical divergences implicating midline DN as well as the CA1 and molecular layer of the HC stood out, especially in modes 1, 3, and 5 of our analysis. The concomitant structural divergences of HC and DN subregions in lonely individuals may in part be explained by the varieties of internally directed cognition that have been consistently associated with each neural system ⍰ such as episodic memory processing, sense of self, and prospective cognition (Addis et al., 2007; Andrews-Hanna et al., 2014b; Hassabis and Maguire, 2009; Szpunar et al., 2014). Especially midline structures of the DN, such as the vmPFC (Campbell et al., 2018), posteromedial cortex (Hebscher et al., 2018; Parvizi et al., 2021), and the RSC (Addis et al., 2007) have been found to serve these types of neural processes. In line with this notion, the fornix ⍰ known to be a key mediator between the HC and midline DN subregions ⍰ has been found to be the most strongly linked to this white-matter fiber in the brain in lonely individuals (Spreng et al., 2020). This observation is of relevance to our present considerations, as direct neuron-to-neuron connections between CA1 and subiculum of the hippocampus with mPFC, OFC, and RSC subregions of the DN have been demonstrated through invasive axonal tracing studies in monkeys (Aggleton et al., 2015; Aggleton et al., 2012; Barbas and Blatt, 1995; Carmichael and Price, 1995; Poletti and Creswell, 1977). Human studies have also found that the functional connectivity strength between the hippocampus and mPFC correlates with the imagination of mental scenes (Campbell et al., 2018; McCormick et al., 2015; Yang et al., 2013), and pre-commissural fornix integrity correlates with episodic richness (Hodgetts et al., 2017; Williams et al., 2020). Indeed, these earlier findings may explain various aspects of the psychological alterations that are characteristic for lonely individuals.

For example, lonely individuals have been described to more frequently reiterate social events from the past, imagine hypothetical or future encounters with others, and reminisce on nostalgic memories (Epley et al., 2008; Epley et al., 2007; Twenge et al., 2003; Wildschut et al., 2006; Zhou et al., 2008). The joint divergences of CA1 and midline DN subregions thus suggest an intimate link between these spatially distributed subregions. A link which a) is supported by the fornix, b) implicates internally directed cognition, and c) is systematically altered in lonely individuals. The notion of CA1, molecular layer, OFC, and mPFC subregions having preferential structural relationships with the fornix receives additional support from our multiple regression and pairwise Pearson’s correlation analyses. Indeed, each of these subregions showed especially strong relationships with the fornix in both analyses. In accord with the growing knowledge of the fornix, it is possible that these concordant findings reflect a difference in the neurocognitive processes necessary for episodic memory retrieval (Tsivilis et al., 2008; Williams et al., 2020). For example, the CA1 and subiculum ⍰ which are the primary origin of pre- commissural fornix projection fibers to the PFC and NAc (Aggleton et al., 2015; Friedman et al., 2002; Jay and Witter, 1991) ⍰ have been found to be crucial for proto-episodic memory in rodents (Buzsáki, 2015; Sun et al., 2020), and detailed episodic autobiographical memory in humans (Bartsch et al., 2011; Zola-Morgan et al., 1986). Hence, the known cognitive upregulation in memory retrieval and social perception biases of lonely individuals may be rooted in how the fornix pathway assists HC-DN communication.

The cognitive biases of the loneliness trait further include perceiving one’s social world as a more threatening place (Cacioppo and Hawkley, 2009; Cacioppo et al., 2016): such as remembering more negative information from past social encounters, and expecting more negative social interactions (Hawkley and Cacioppo, 2010). Based on these earlier psychological insights, we expected that the subregions with multiple divergences in lonely individuals would show a strong relationship to the NAc, a key node of the brain’s reward circuitry that has previously been associated with loneliness (Kiesow et al., 2020). In support of this idea, our multiple regression analysis revealed that CA1, molecular layer, vmPFC and OFC each showed strong relationships with the NAc. The seemingly separate psychological and behavioral characteristics of loneliness may thus be a product of the remodeling of integrated neural systems. This notion receives further support from the similarity in the subregion associations revealed through our separate multiple regression models of the hippocampus. In these, we observed a strong overlap between the hippocampal subregions having strong relationships with either the NAc or fornix. Indeed, the NAc receives the bulk of its hippocampal inputs through the pre-commissural fornix pathway (Kelley and Domesick, 1982; Poletti and Creswell, 1977). Hence, our results substantiate previous findings on cross-associations between the CA1 of the hippocampus, fornix, mPFC, and NAc.

In addition to the collective divergences in hippocampal and mPFC subregions in lonely individuals, our analysis pinpointed parallel findings in additional DN subregions. In particular, the majority of DN subregion divergences we found were in the PCC, and especially RSC. In general, these subregions also showed strong structural relationships with the fornix in our regression analyses. In line with our results from brain-imaging modelling, direct axon tracing studies in rodents have documented CA1 and subiculum as the main hippocampal projection sites to the RSC (Cenquizca and Swanson, 2007; Wyass and Van Groen, 1992). Monkey experiments have additionally reported that direct HC-RSC projections are chiefly mediated through the subiculum – a close interaction partner of hippocampal CA1 (Aggleton et al., 2012; Kobayashi and Amaral, 2003; Rosene and Van Hoesen, 1977; Wyass and Van Groen, 1992). In conjunction with the CA1 structural divergences we observed in lonely individuals, we also observed 4 total subiculum divergences across all examined signatures. Hence, our population neuroscience evidence for inter-dependence between CA1 and subiculum in the allocortex and RSC subregions in the neocortex directly confirms established knowledge from invasive animal experiments (Nitzan et al., 2020). Thus, an attractive interpretation of these findings of structural divergences in RSC subregions for lonely individuals (in modes 1, 5, 9, 14, and 15) invokes a difference in the facility of visualizing internally centered thoughts. Indeed, amongst the scant knowledge of the function of the human RSC, this part of the posterior midline is believed to assist in spatial navigation, episodic memory, and visual details of mental scene construction (Andrews-Hanna et al., 2014a; Hassabis and Maguire, 2009; Mitchell et al., 2018; Morton et al., 2021; Vann et al., 2009).

Moreover, RSC-mediated mental scene construction could be one aspect of a more general role the DN has in spontaneous cognition ⍰ such as task-free ‘mind-wandering’ (Dohmatob et al., 2020b; Smallwood et al., 2021). In fact, the hippocampus has been found to support stimulus-independent neural activity (Pastalkova et al., 2008), which would be a perfect partner for these DN functions, via so-called sharp wave ripples (SWR) (Buzsáki, 1986; Diba and Buzsáki, 2007). These phenomena are manifestations of spontaneous neural activity synchronizations in the hippocampus circuit that occur independent of any environmental cues, or externally structured tasks (Buzsáki and Vanderwolf, 1983; Ylinen et al., 1995). SWR have been carefully studied through electrophysiological experiments in rodents and other animals, as opposed to most of the neural activity specific to regions of the human DN (Buzsáki, 2015; Csicsvari et al., 2000). Of relevance for our present considerations, CA1 pyramidal neurons are purported to underlie the expansive ‘ripples’ of SWR activity. The preceding ‘sharp wave’ on the other hand is most pronounced in the apical dendrites of CA1 neurons (Buzsáki, 2015). Along this line, the hippocampus atlas used for our study collapsed the apical dendrites (both the stratum radiatum and stratum lacunosum moleculare) of subiculum and CA neurons into one composite subregion: the molecular layer (Iglesias et al., 2015). Thus, the molecular layer and CA1 of our set of human hippocampus subregions offered a way to analyze systematic alterations of the neural substrates shown to be highly visible locations of SWR activity in the hippocampus of rodents and monkeys.

Indeed, the CA1 and molecular layer had the greatest number of total hits in lonely individuals. More specifically, there were 7 total hits in the molecular layer and 6 total hits in CA1 across all examined modes. Since our analyses revealed co-occurring divergences in midline DN subregions, SWR activity suggests itself as a compelling interpretation that can accommodate these collective results. The sensory-distal processing regimes realized by the DN may be perfectly attuned to integrating spontaneous SWR activity. For example, in rodents, CA1 cell activity and coupled SWR have been shown to assist in prospective cognitive processes (Pfeiffer and Foster, 2013), such as simulating spatial trajectories that were never experienced before (Dragoi and Tonegawa, 2011; Gupta et al., 2010; Ólafsdóttir et al., 2015). A functional interaction between hippocampal SWR activity and association cortex regions, such as the RSC, has even been demonstrated in animal species (Khodagholy et al., 2017; Nitzan et al., 2020). For example, in rats, it has been shown that auditory cortex activity predicts subsequent SWR activity (Rothschild et al., 2017). Conversely, hippocampal activity also predicted subsequent activity in the auditory cortex. Based on these results, the authors proposed that immediately before and after SWR occur there is rapid information flow in a cortical–hippocampal–cortical loop (Rothschild et al., 2017). Similarly, studies in primates (Kaplan et al., 2016; Logothetis et al., 2012) have also identified the RSC and PCC to be the regions of the neocortical mantle that are most closely coupled with hippocampal SWR (Logothetis et al., 2012). Our findings of selective structural divergences in both the CA1 and molecular layer as well RSC subregions across modes is thus very suggestive of a difference in the allocortical-neocortical information processing pathways that involve SWR activity.

Facilitation of mental scene construction is discussed as one of the main functional contributions of the hippocampus (Dohmatob et al., 2020b; Hassabis and Maguire, 2009). It thus appears plausible that a systematic alteration of SWR activity may affect the vividness of an individual’s episodic memories and imaginings (Norman et al., 2019). In fact, SWR activity has been proposed to underlie trajectory sequence replay in rodents and as well as ‘offline’ states of human cognition, such as daydreaming (György Buzsáki, 2019). Thus, the increased frequency and intensity of nostalgic reminiscence which has been associated with loneliness may potentially also be associated with altered SWR activity (Zhou et al., 2008). A strong relationship between SWR and loneliness is additionally suggested by their shared association with reward processing. Indeed, loneliness alters the subjectively perceived valence of social and non-social reward in humans (Cacioppo et al., 2009), while SWR activity has been demonstrated to be responsive to reward-contingent processing in animals (Ambrose et al., 2016; Singer and Frank, 2009; Sosa et al., 2020). Similarly, the fundamental place cell representations of the CA1 pyramidal cells, which underlie SWR activity (cf. above), were reported to intimately relate to reward processing (De Lavilléon et al., 2015; Robinson et al., 2020). Certain populations of CA1 cells have even been found to act as a dedicated channel for reward-related processing across environments (Gauthier and Tank, 2018). These findings and conclusions from previous animal experiments therefore appear to dovetail with our analyses in people highlighting the NAc with CA1 and molecular layer. We hence wonder whether the attempt to fill the perceived social void through mental imagery in lonely individuals coincides with interlocked structural alterations in the allocortical-neocortical co-variations.

However, the relationship between the structure of the HC and DN with the tendency to feel socially disconnected from others may only be partially determined by one’s environment and life experience. Indeed, our genome-wide analyses showed that participant-specific expressions of HC-DN co-variation have diverging links to the heritable components of loneliness and low social support. This is in accord with previous research showing that loneliness has consistent, yet subtle genetic underpinnings (Spithoven et al., 2019). For example, twin studies have identified genetic contributions to individual differences in feelings of loneliness that is as high as ∼48% (Boomsma et al., 2005). Recent genome-wide association studies with large sample sizes (e.g., n= 452,302 (Day et al., 2018)) have also reported the contribution of common genetic variants to loneliness to range from 4-27% (Abdellaoui et al., 2019a; Day et al., 2018; Gao et al., 2017). These earlier genome-wide analyses have additionally pointed out a small number of specific gene loci that are significantly associated with loneliness and regular participation in social groups (Day et al., 2018).

An underlying genetic contribution to the experience of loneliness is further supported by recent genome-wide correlation analyses. For example, one study has demonstrated that our UK Biobank loneliness trait significantly shared underlying genetic factors with each of 264 different demographic, lifestyle and disease phenotypes (Spreng et al., 2020). Another study has demonstrated a strong genetic correlation between loneliness and neuroticism (Abdellaoui et al., 2019a). Taken together, these recent population neuroscience studies indicate that the genetic determinants underlying loneliness are probably quite polygenic and involve complex gene-environment interactions. In our study, these previous insights were extended by relating PRS for loneliness with the inter-individual expression of multiple spatially overlapping signatures of HC-DN co-variation. Importantly, we have identified a select subset of HC-DN co- variation modes that had a significant relationship with genetic predisposition for loneliness.

In particular, we found a significant heritability effect for the expression of the loneliness specific signature in modes 8 and 9 ⍰ both of which also showed multiple subregion divergences in lonely individuals. We also noted a particular concordance between our PRS analysis of loneliness with our uncovered subregion-specific divergences in mode 9. Intriguingly, in our multiple regression analyses of the NAc, each subregion which showed divergence in mode 9 for loneliness (i.e., OFC, vmPFC, pSTS, and left parasubiculum) were found to have a strong relationship with the NAc. These convergent results may suggest that a specific aspect of the genetic predisposition for loneliness consists in innate tendencies involving reward-related processing. This view would be in line with past findings associating loneliness with social reward valence (Cacioppo et al., 2009), substance abuse or dependence (Abdellaoui et al., 2019b; Åkerlind et al., 1992), executive function (Baumeister et al., 2005; Hawkley et al., 2009), and impulsive behaviors (Layden et al., 2017). Recently, it has even been reported that social isolation is linked to altered neural activity responses in midbrain regions and induces social craving in a way that may be similar to how fasting causes hunger (Tomova et al., 2020). However, our findings on how structural HC-DN co-variation relates to the genetic predisposition for social isolation may be better framed in terms of susceptibility to the environment.

Such as the previously proposed notion that the driving forces behind genetic heritability are not direct, but rather come into effect through an altered sensitivity to environmental conditions (Belsky et al., 2007). For example, while oxytocin and social support have been shown to interact in the stress response (Heinrichs et al., 2003), a single nucleotide polymorphism in the oxytocin receptor gene (OXTR) has been found to differentially affect stress response depending on adequate social interactions with others (Chen et al., 2011). Indeed, the heightened levels of stress in humans with social isolation has been argued to be fundamentally different from a simple general and diffuse stress response (Cacioppo and Cacioppo, 2018b; Zayan, 1991).

In fact, animal studies have afforded detailed accounts of the effects of social isolation stress on the hippocampus (Sapolsky, 1996), focusing in on the apical dendrites of CA neurons (Conrad et al., 2017). When considered in conjunction with this animal literature, our analysis could therefore extend our incomplete understanding of the brain basis of social isolation-related stress by identifying the precise allocortical and neocortical anatomical subregions at play. A possible manifestation of the effects of social isolation-related stress in our study is the pronounced total number of divergences in the molecular layer. In particular, we found 7 total hits in the molecular layer across all examined modes for lonely individuals. As mentioned previously, this molecular layer subregion consisted of a combination of the apical dendrites of subiculum and CA neurons ⍰ with CA1 composing the largest portion (Iglesias et al., 2015). In light of this, the observation of preferential hits in the molecular layer of lonely individuals broadly aligns with past studies on chronic stress in rodents, tree shrews, and non-human primates. Indeed, experimental studies in these species have reported detailed cellular and physiological consequences of induced stress on the hippocampus and other related brain regions.

In the hippocampus specifically, it has been shown that chronic stress results in the selective atrophy of the apical dendrites of CA3 cells, but not CA3 basal dendrites (Magarin and McEwen, 1995; Magariños et al., 1997; Magariños et al., 1996; Watanabe et al., 1992; Woolley et al., 1990). In contrast, elevated chronic stress in the face of multiple stressors results in additional atrophy of the apical dendrites of CA1 cells, yet not CA1 basal dendrites (Alfarez et al., 2008; Maras et al., 2014). Based on these findings, researchers have proposed that CA3 apical dendrites are the most susceptible hippocampal structure to chronic stress. Yet, stress affects CA1 apical dendrites only with a more severe experimental stressor than that needed to affect CA3 apical dendrites (Conrad et al., 2017). Our findings of preferential hits in the molecular layer of lonely individuals may thus be a marker of elevated stress, and relate to the physiological characterization of loneliness as a trait coinciding with elevated stress and immune response levels (Adam et al., 2006; Cacioppo et al., 2000; Cole and immunity, 2008; Hawkley and Cacioppo, 2010; Steptoe et al., 2004; Xia and Li, 2018).

Building upon this, our concerted analysis framework for HC and DN subregion variation thus allowed us to relate the potential effects of chronic stress on neocortical partners in the DN. For example, in mode 1 we observed 4 hits in the molecular layer with coincident hits in midline DN subregions, such as the dmPFC, PCC, and RSC. The collection of parallel findings of divergences between the stress-susceptible molecular layer in the allocortex and specific midline cortical subregions might therefore reflect the effects of stress on the higher associative cortex.

Indeed, previous studies have shown that the effects of chronic stress on the hippocampus percolate into interaction partners with known axonal connections, such as target areas in the mPFC (Aggleton et al., 2015; Barbas and Blatt, 1995; Carmichael and Price, 1995; Cerqueira et al., 2007; Yuan et al., 2021). For example, in one study where rats were submitted to 4 weeks of chronic stress, they were found to have a deficit in hippocampal-PFC synaptic strength compared to controls (Cerqueira et al., 2007). It was also found that the stressed rats had lower PFC volume, and poorer performance on working memory and behavioral flexibility tasks ⍰ activities which are thought to depend on mPFC activity (Cerqueira et al., 2007). Another more recent study has demonstrated that amongst squirrel monkeys with bouts of pre-pubertal social isolation, there was an association between functional hyper-connectivity in PFC–subcortical circuits (e.g., dmPFC, vmPFC, OFC; amygdala, ventral striatum, HC) with reduced anxiety-like behavior in later adulthood. In line with these previous experiments, our analytical strategy in the UK Biobank population allowed us to identify concomitant stress-related divergences in the molecular layer of the hippocampus and divergences in the mPFC, PCC, and RSC of the human DN.

It is well-known that chronic stress causes measurable neural consequences for the hippocampus in different animal species (Champagne et al., 2008; Conrad, 2008; McEWEN and Magarinos, 1997). Yet, the impact of social isolation and chronic stress on the hippocampus has repeatedly been argued to be reversible in a matter of a few weeks with social rehabilitation, or reintroduction to an enriched environment (Biggio et al., 2019; Hutchinson et al., 2012; Ibi et al., 2008; Sandi et al., 2003). In a similar vein, in the higher association cortex, the effects of chronic stress on PFC dendritic morphology in rats and functional connectivity in humans have been shown to be reversible after stressful experiences (Bloss et al., 2010; Liston et al., 2009; Radley et al., 2005; Soares et al., 2012). In light of this past literature, there is likely to be elasticity to the effects of social isolation and perhaps the stress-related volumetric divergences that we uncovered in the hippocampus and its neocortical interaction partners. Indeed, social rehabilitation by returning to the usual social environment may remold the examined allocortical-neocortical brain circuits.

## Acknowledgements

We are grateful to Adrien Peyrache and David Redish for useful comments on a previous version of the manuscript.

This project has been made possible by the Brain Canada Foundation, through the Canada Brain Research Fund, as well as by NIH grant R01AG068563A and the Canadian Institutes of Health Research. DB was also supported by the Healthy Brains Healthy Lives initiative (Canada First Research Excellence fund), and by the CIFAR Artificial Intelligence Chairs program (Canada Institute for Advanced Research), as well as Research Award and Teaching Award by Google.

## Author contributions

CZ, RNS, DB conceived and designed research; CZ and DB performed experiments; CZ and DB analyzed data; CZ and DB interpreted results of experiments; CZ and DB prepared figures; CZ and DB drafted manuscript; CZ, RNS, and DB edited and revised manuscript; CZ, RNS, and DB approved final version of manuscript

