## Supplementary Online Material for "Loneliness is linked to specific subregional alterations in hippocampus-default network co-variation"

**Supplementary online mat****erial**

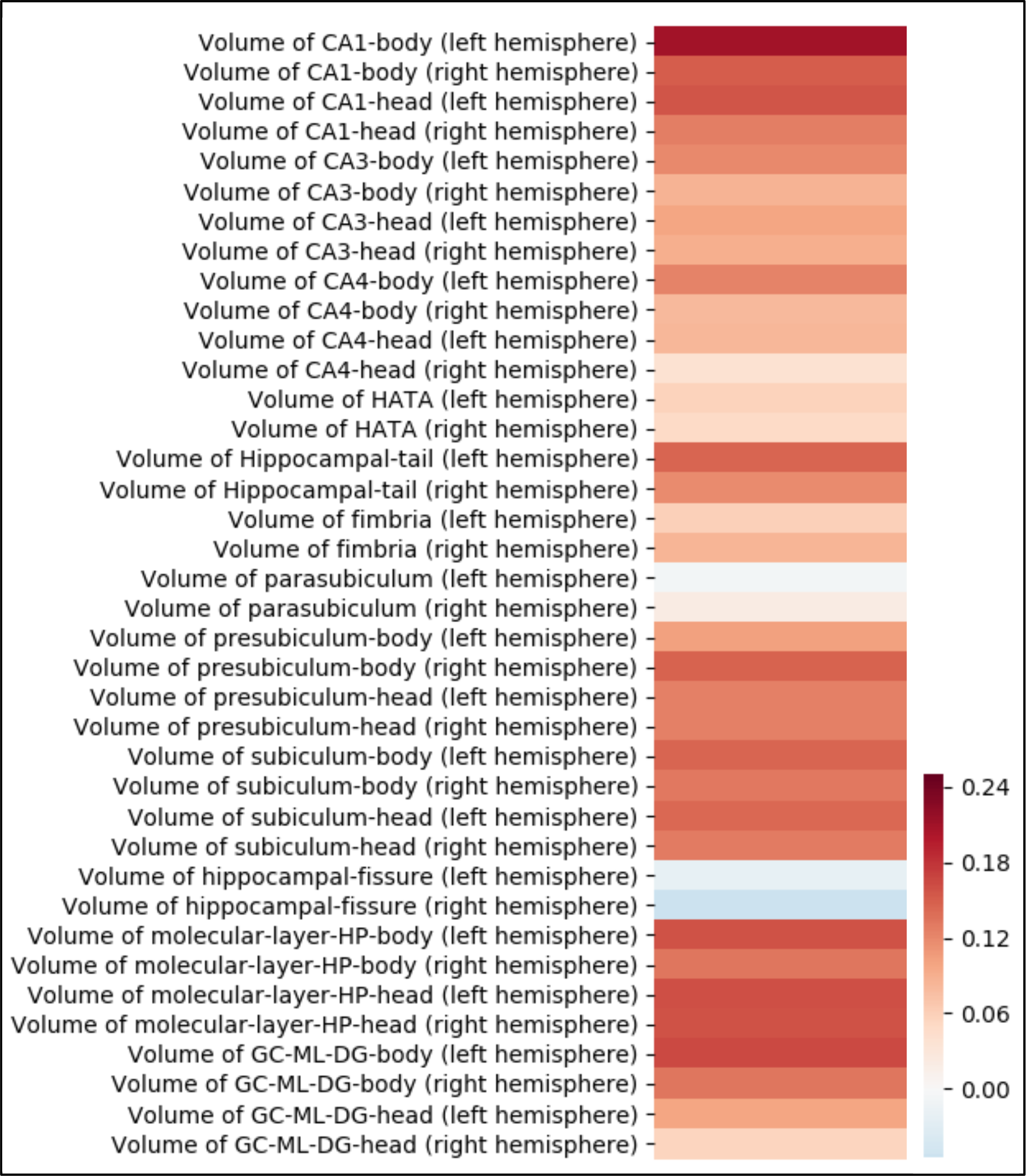

**Supplementary Figure 1. Simple pairwise correlation between fornix microstructural integrity and hippocampal subregion volumes.** Pearson correlation coefficients were computed for each hippocampal subregion with the microstructural integrity of the fornix white-matter tract. This analysis was based on ~38,701 participants in the UK Biobank dataset. Hot colors depict a positive correlation between a hippocampal subregion and fornix structure. Cold colors indicate a negative correlation. The volume of CA1 body of left hippocampus showed the strongest relationship with fornix structure. HATA = hippocampal amygdala transition area, GC-ML-DG = granule cell layer of dentate gyrus.

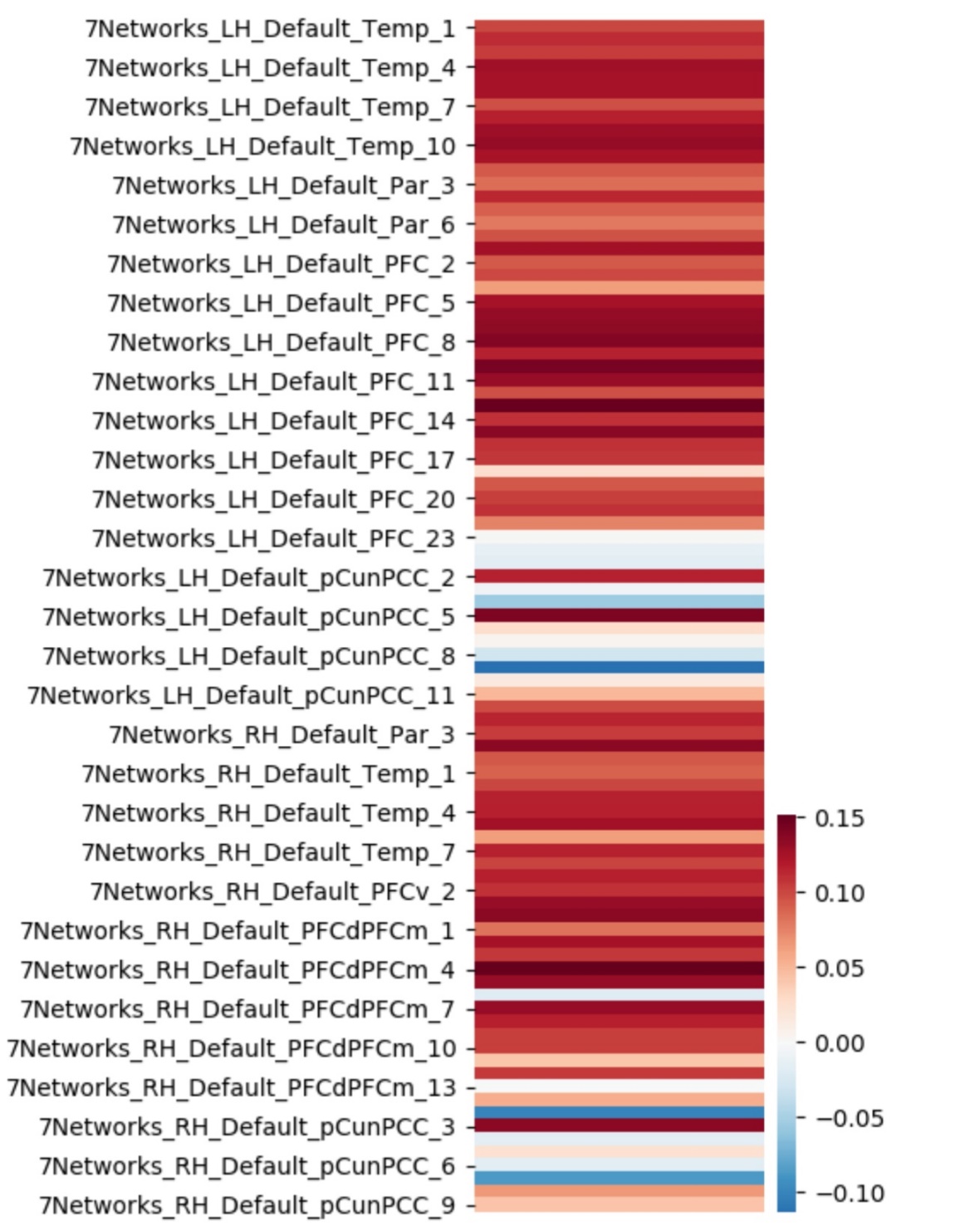

**Supplementary Figure 2. Simple pairwise correlation between fornix microstructural integrity and default-network subregion volumes.** Pearson correlation coefficients were computed for each default-network subregion with the microstructural integrity of the fornix white-matter tract. This analysis was based on ~38,701 participants in the UK Biobank dataset. The volume of bilateral posterior cingulate and retrosplenial cortex subregions showed, in general, the weakest or strongest anti-correlation with fornix structure, when compared to prefrontal, temporal, and parietal subregions. Temp = temporal lobe region, Par = parietal lobe region, PFC = prefrontal cortex region, PCC = posterior cingulate cortex region, LH = left hemisphere, RH = right hemisphere.

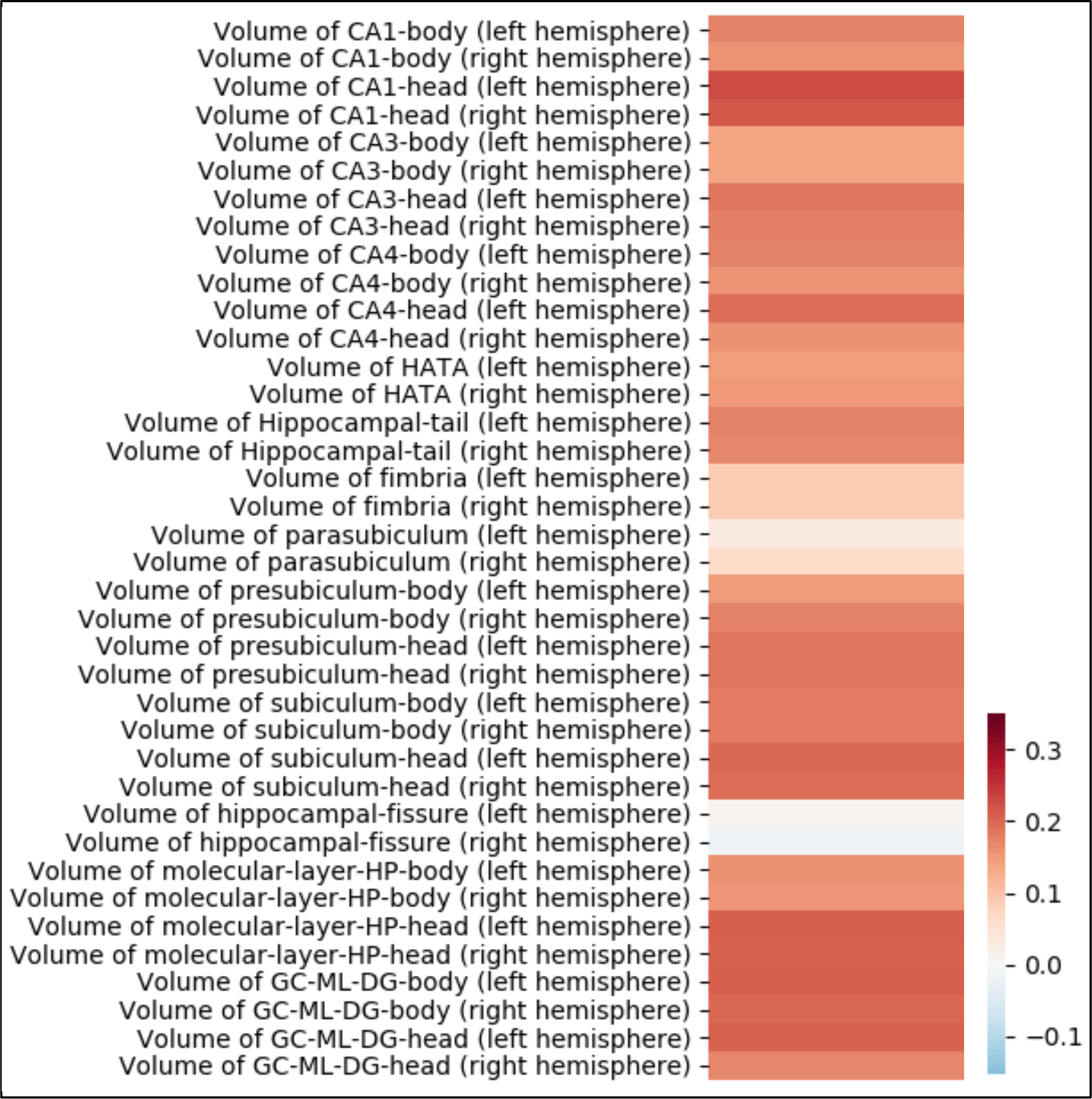

**Supplementary Figure 3. Simple pairwise correlation between nucleus accumbens and hippocampal subregion volumes.** Pearson correlation coefficients were computed for each hippocampal subregion with the volume of the mean nucleus accumbens volume (left and right hemispheres). This analysis was based on ~38,701 participants in the UK Biobank dataset. The volume of bilateral CA1 body head showed the strongest relationship with nucleus accumbens volume. HATA = hippocampal amygdala transition area, GC-ML-DG = granule cell layer of dentate gyrus.

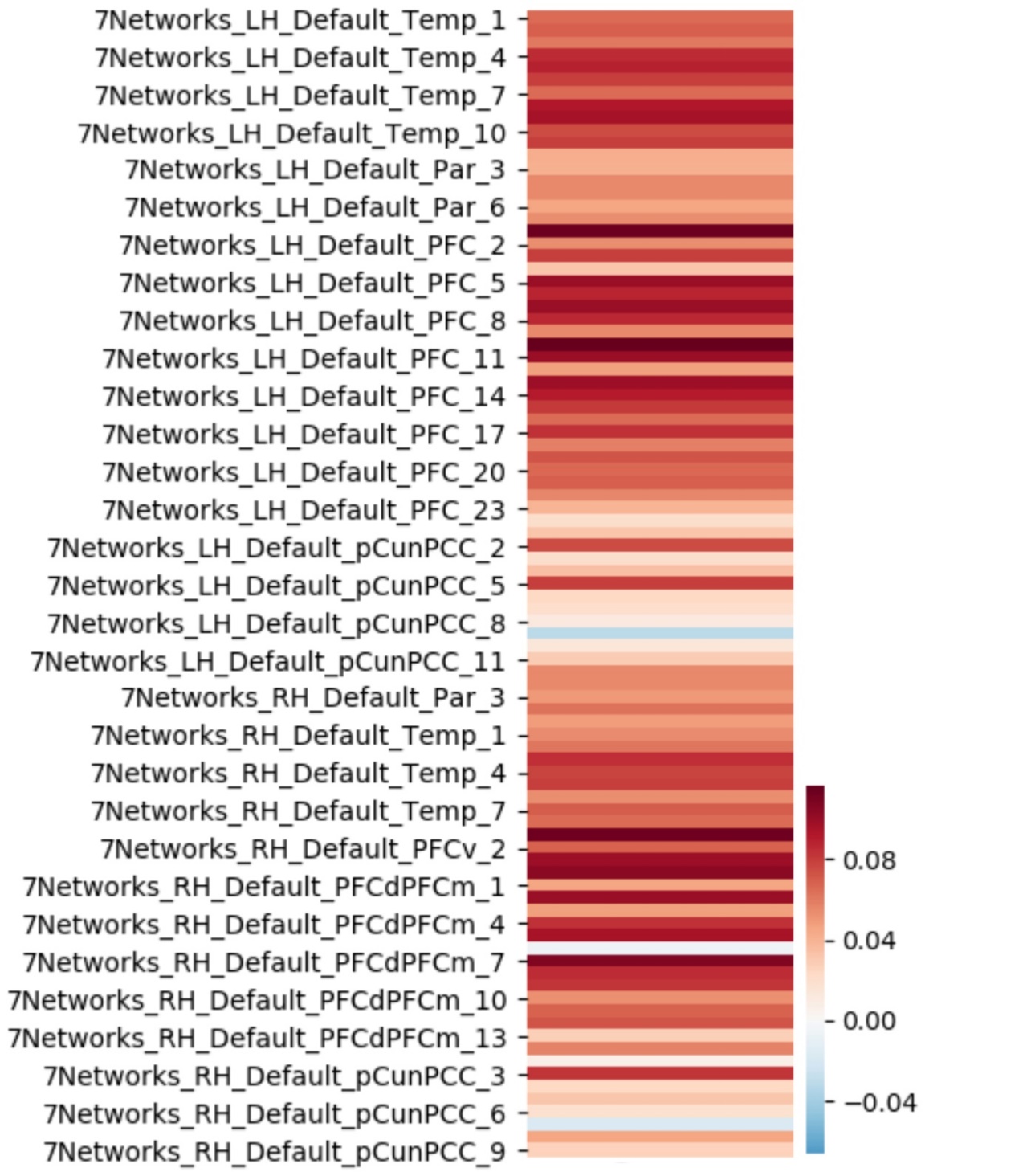

**Supplementary Figure 4. Simple pairwise correlation between nucleus accumbens and hippocampal subregion volumes.** Pearson correlation coefficients were computed for each default-network subregion with the mean nucleus accumbens volume (left + right hemispheres). This analysis was based on ~38,701 participants in the UK Biobank dataset. The volume of bilateral orbitofrontal cortex had the strongest relationship with nucleus accumbens volume. Conversely, bilateral posterior cingulate and retrosplenial cortex subregions showed the weakest independent relationships with nucleus accumbens volume. Temp = temporal lobe region, Par = parietal lobe region, PFC = prefrontal cortex region, PCC = posterior cingulate cortex region, LH = left hemisphere, RH = right hemisphere.

| **Mode** | **Explained Variance (rho * 100)** |
| --- | --- |
| 1 | 51.0 |
| 2 | 42.2 |
| 3 | 38.9 |
| 4 | 30.9 |
| 5 | 26.8 |
| 6 | 23.3 |
| 7 | 22.2 |
| 8 | 20.1 |
| 9 | 17.8 |
| 10 | 16.8 |
| 11 | 15.0 |
| 12 | 14.7 |
| 13 | 13.8 |
| 14 | 11.9 |
| 15 | 11.5 |
| 16 | 11.1 |
| 17 | 9.9 |
| 18 | 9.2 |
| 19 | 8.4 |
| 20 | 7.9 |
| 21 | 7.4 |
| 22 | 7.2 |
| 23 | 6.6 |
| 24 | 6.2 |
| 25 | 6.1 |

**Supplementary Table 1. Relative explained variance of each of our 25 candidate hippocampus-default network signatures.** The canonical correlation (as Pearson’s rho * 100) of each of our 25 candidate modes identified through a canonical correlation analysis of hippocampus and default network subregion volumes.

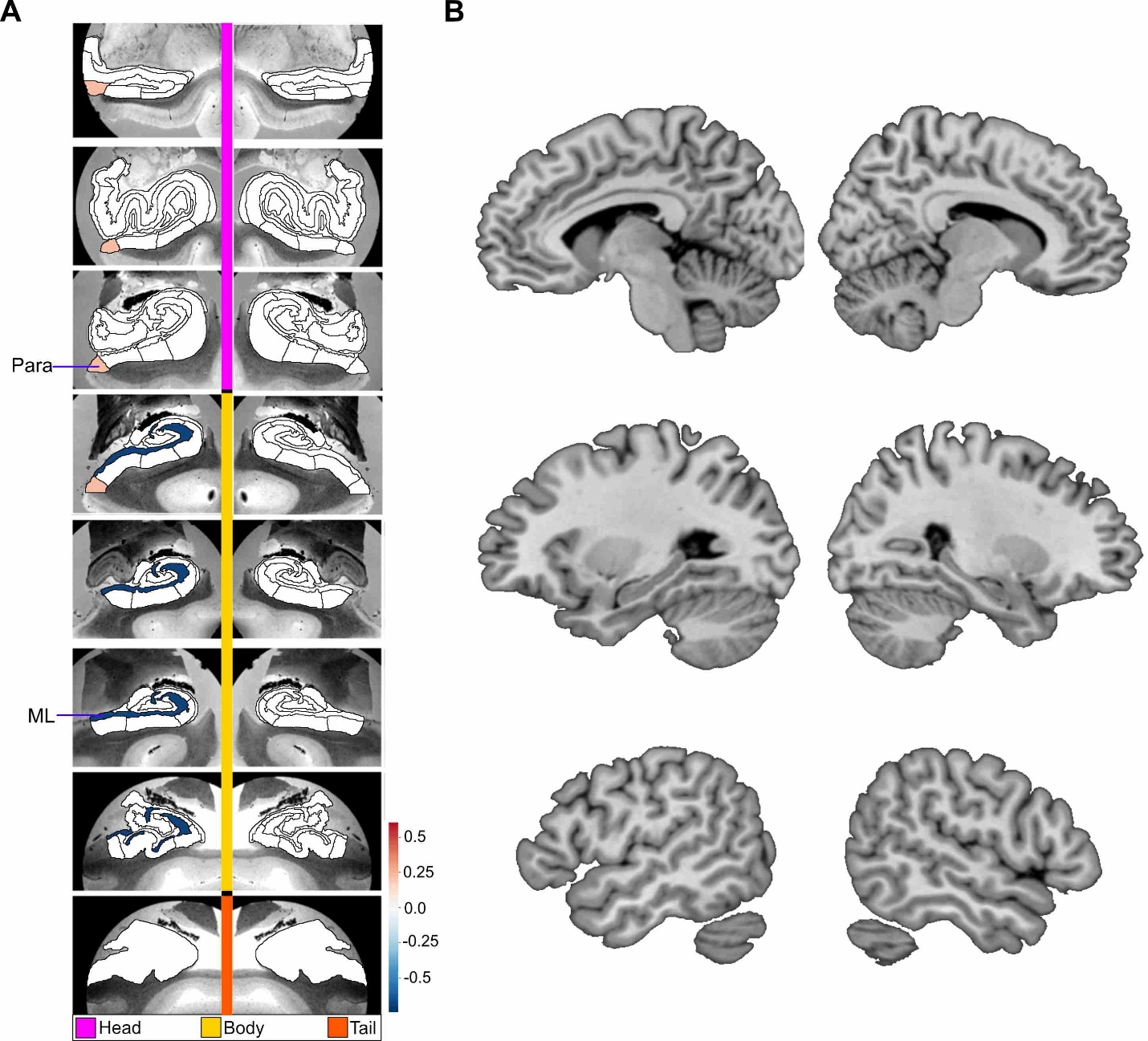

**Supplemental Figure 5. Loneliness is associated with structural divergences preferentially in left parasubiculum and left molecular layer within mode 2 of hippocampus-default network co-variation.** Shown here are the subregion divergences in mode 2. Mode 2 of the CCA solution achieves the second most explanatory hippocampal-default network co-variation signature, with a canonical correlation of rho = 0.42. **A** exhibits the hippocampal (HC) subregion patterns (left, one cannical vector of mode 2) that robustly diverge between lonely and non-lonely groups in mode 2. **B** shows the default network (DN) subregion patterns (right, other canonical vector of mode 2) of the DN (right) that robustly diverge between lonely and non-lonely groups. ML = molecular layer, Para = parasubiculum.

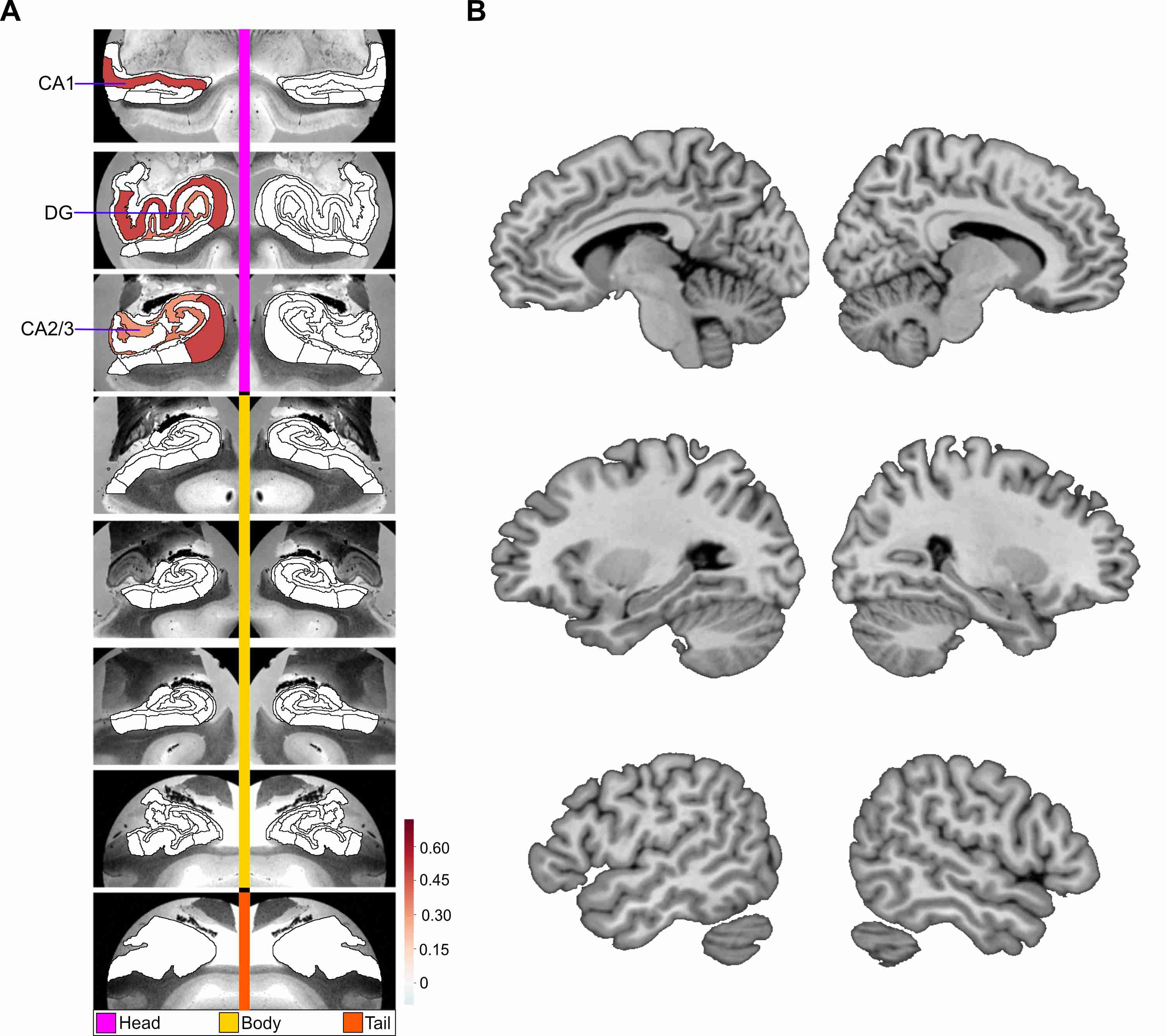

**Supplemental Figure 6. Loneliness is associated with preferential divergences in the head portion of the traditional hippocampal ‘tri-synaptic circuit’ within mode 8 of hippocampus-default network structural co-variation.** Shown here are the subregion divergences in mode 8. Mode 8 of the CCA solution achieves the eight most explanatory hippocampal-default network co-variation signature, with a canonical correlation of rho = 0.20. **A** exhibits the hippocampal (HC) subregion patterns (left, one cannical vector of mode 8) with paramater weights that robustly diverge between lonely and non-lonely groups in mode 8. **B** shows the default network (DN) subregion patterns (right, other canonical vector of mode 8) of the DN (right) that robustly diverge between lonely and non-lonely groups. The trisynaptic circuit is a well recognized flow of neural information in the hippocampus, with synaptic connection from dentate gyrus onto to CA3, and then CA3 onto CA1. Overall, the divergences in mode 8 are the subregions of the tri-synaptic circuit, but only in the left hemisphere.

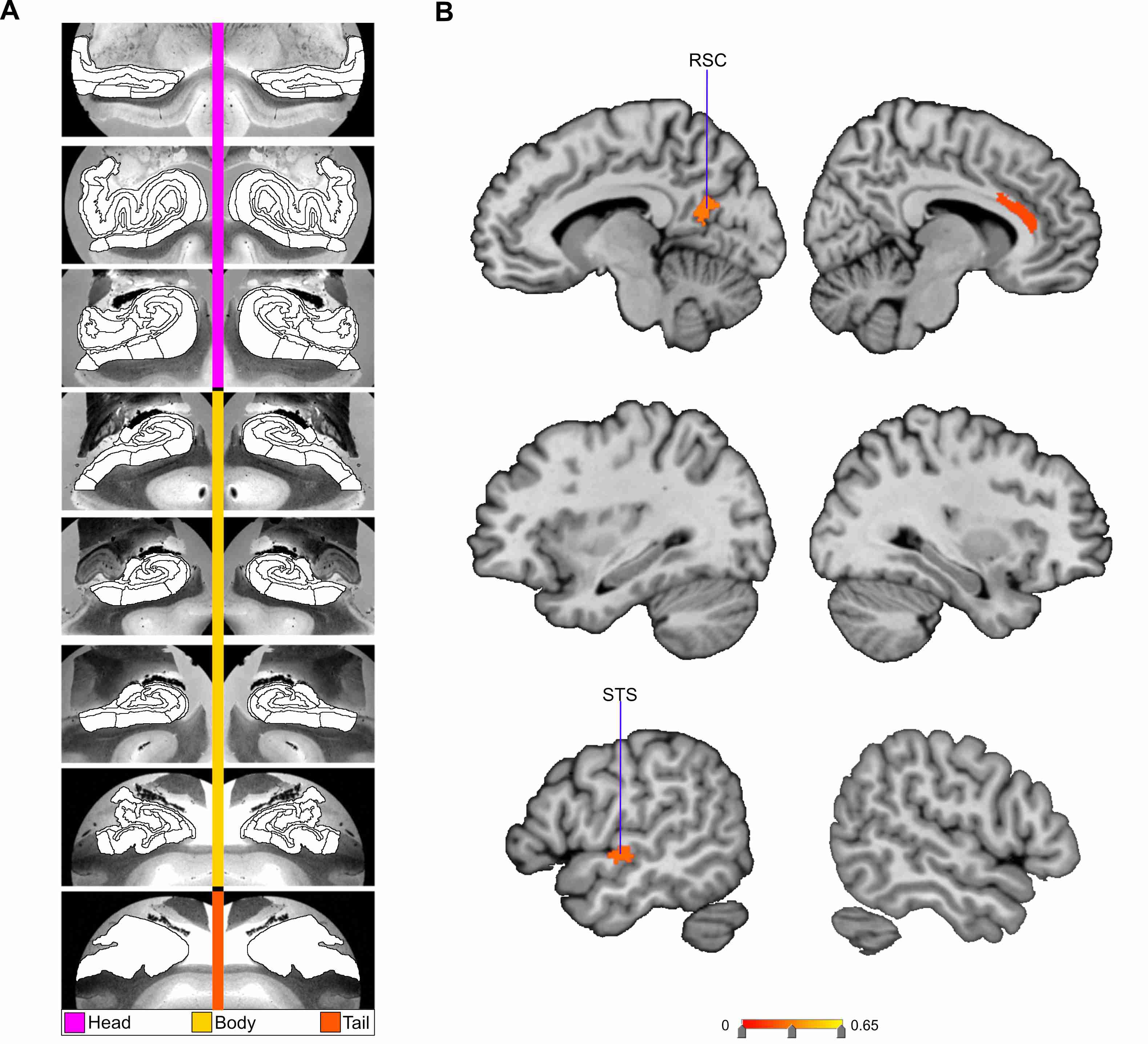
**Supplemental Figure 7. Loneliness is associated with divergences in DN subregions within the mode 14 of hippocampus-default network structural co-variation.** Shown here are the subregion divergences in mode 14 of hippocampus-DN covariation. Mode 14 of the CCA solution achieves the fourteenth most explanatory hippocampal-default network co-variation signatures, with a canonical correlation of rho = 0.12. **A** exhibits the hippocampal (HC) subregion patterns (left, one cannical vector of mode 14) with paramater weights that robustly diverge between lonely and non-lonely groups. **B** shows the default network (DN) subregion patterns (right, other canonical vector of mode 14) of the DN (right) that robustly diverged between lonely and non-lonely groups. Overall, there are divergences for in left retrosplenial cortex, right anterior cingulate cortex, and left superior temporal gyrus in lonely individuals.

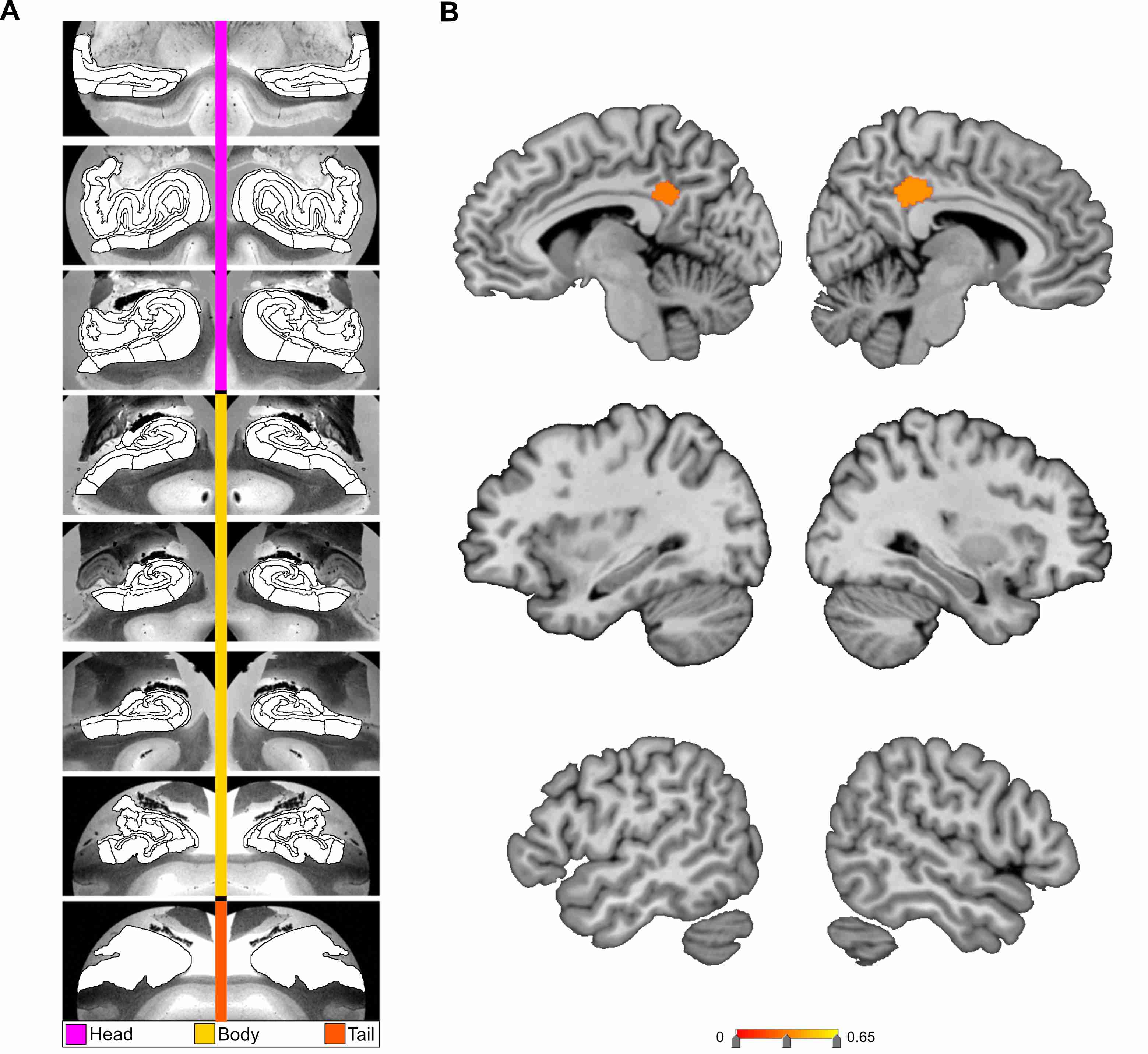

**Supplemental Figure 8. Loneliness is associated with bilateral divergences in posterior midline subregions of the DN within mode 15 of hippocampus-default network structural co-variation.** Shown here are the subregion divergences in mode 15 of hippocampus-DN covariation. Mode 15 of the CCA solution achieves the fifteenth most explanatory hippocampal-default network co-variation signatures, with a canonical correlation of rho = 0.12. **A** exhibits the hippocampal (HC) subregion patterns (left, one cannical vector of mode 15) with paramater weights that robustly diverge between lonely and non-lonely groups. **B** shows the default network (DN) subregion patterns (right, other canonical vector of mode 15) that robustly diverged in the posterior cingulate cortex between lonely and non-lonely groups. Overall, there are divergences in lonely individuals in bilateral posterior cingulate subregions in mode 15.
